# Novel Apelin-expressing gCap Endothelial Stem-like Cells Orchestrate Lung Microvascular Repair

**DOI:** 10.1101/2021.07.12.452061

**Authors:** Rafael Soares Godoy, David P Cook, Nicholas D Cober, Emma McCourt, Yupu Deng, Liyuan Wang, Kenny Schlosser, Katelynn Rowe, Duncan J Stewart

**Author notes:** **Corresponding author**: Duncan J Stewart., Executive Vice-President Research, The Ottawa Hospital; CEO & Scientific Director, Ottawa Hospital Research Institute; The Evelyne & Rowell Laishley Chair and Professor, Department of Medicine, University of Ottawa, 501 Smyth Road, Box/C.P. 511, Ottawa, ON, K1H 8L6.

## Abstract

**Question:** We sought to define the mechanism underlying lung microvascular regeneration in a severe acute lung injury (ALI) model induced by selective lung endothelial cell ablation.

**Methods:** Changes in lung cell populations and gene expression profiles were determined in transgenic mice expressing human diphtheria toxin (DT) receptor targeted to ECs using single-cell RNA sequencing at baseline (day 0) and days 3, 5 and 7 after lung EC ablation.

**Results:** Eight distinct endothelial clusters were resolved, including alveolar aerocytes (aCap) ECs expressing apelin at baseline, and general capillary (gCap) ECs expressing the apelin receptor. Intratracheal instillation of DT resulted in ablation of >70% of lung ECs, producing severe ALI with near complete resolution by 7 days. At 3 days post injury, a novel gCap population emerged characterized by *de novo* expression of apelin, together with the stem cell marker, protein C receptor. These stem-like cells transitioned to proliferative ECs, expressing apelin receptor together with the pro-proliferative transcription factor, *FoxM1*. This progenitor-like cell population was responsible for the rapid replenishment of all depleted EC populations by 7 days post injury, including aerocytes which play a critical role in re-establishment of the air-blood barrier. Treatment with an apelin receptor antagonist prevented recovery and resulted in excessive mortality, consistent with a central role for apelin signaling in EC regeneration and microvascular repair.

**Conclusion:** The lung has a remarkable capacity for microvasculature EC regeneration which is orchestrated by signaling between newly emergent apelin-expressing gCap endothelial stem-like cells and highly proliferative, apelin receptor positive endothelial progenitors.

**Take-Home message:** Using sublethal lung endothelial cell (EC) ablation, we show for the first that EC regeneration and resolution of acute lung injury is orchestrated by novel apelin-expressing, gCap endothelial stem-like cells by a mechanism requiring apelin signaling.

**Graphical Abstract:** 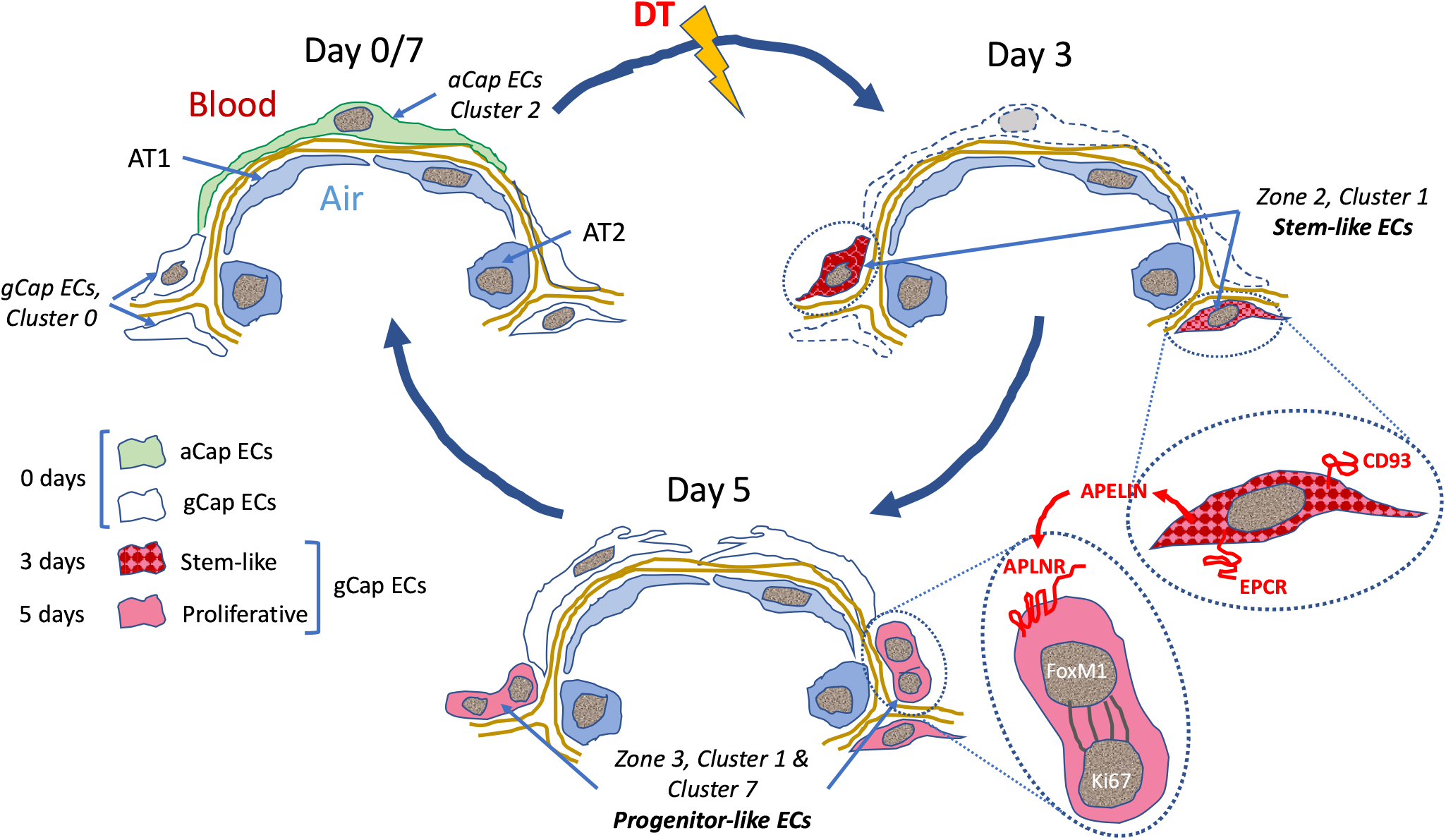

A schematic representation of EC populations contributing to microvascular repair. At baseline (Day 0), there are two main alveolar groups of capillary ECs: larger apelin positive aCap ECs, termed aerocytes, that play a key structural role in forming the air-blood barrier; and smaller apelin receptor (*Aplnr*) expressing gCap ECs, which are found in the thicker regions at the corners of the alveoli. After DT-induced EC ablation, there is a marked depletion of both EC populations and the appearance of novel transitional and transient populations. At Day 3, there is the appearance of stem-like gCap ECs that paradoxically express apelin, but not its receptor, and are characterized by various stem and progenitor cell markers but show no evidence of proliferation. By Day 5, these transition to ECs expressing *Aplnr* which have a strong proliferative phenotype, as evidenced by FoxM1 and Ki67 expression, and then rapidly replenish depleted EC pools, including aCap ECs, by Day 7. This transition is orchestrated by the interaction of apelin with its receptor as a critical mechanism in lung microvascular regeneration after EC injury. AT1 = alveolar type −1 epithelial cell; AT2 = alveolar type-2 epithelial cell; APLNR = apelin receptor; ANGPT2 = angiopoietin 2; EPCR = Endothelial protein C receptor.

## Introduction

Acute lung injury (ALI) and its severe clinical counterpart, the adult respiratory distress syndrome (ARDS), remains a major cause of morbidity and mortality in critically ill patients, accounting for ~30% of ICU admission with a 28-day mortality approaching 40%(1,2). Despite decades of research, no specific therapies have been developed that improve outcomes in ARDS(1). The COVID-19 pandemic has highlighted the devastating nature of this condition as infection with the SARS CoV2 virus leads to a particularly severe form of ARDS(3,4) which has been responsible for the vast majority of over 5 million deaths world-wide. COVID-19 associated ARDS has a strong vascular component which is characterized by intense endothelial inflammation (endothelialitis) and necrosis,(5,6) consistent with the emerging role of endothelial injury in other forms of ALI leading to breakdown of the air-blood barrier(7). Indeed, recent reports have suggested that endothelial repair is required for resolution of ALI(8), and thus an important target for the development of novel therapeutic strategies. Unfortunately, little is known about the mechanisms that underlie lung macrovascular repair and its role in ALI resolution. In experimental models the transcription factor, Forkhead box M1 (*FoxM1*), has been implicated as a driver of EC proliferation and microvascular repair that is required for ALI resolution(9–11), and apelin has been reported to protect against inflammation and oxidative stress(12). During angiogenesis, apelin is induced in endothelial tip cells by tissue hypoxia and VEGF and signals to trailing stalk ECs that express the apelin receptor(13,14). Even though apelin is known to play an important role in vascular development and angiogenesis,(14,15) the relevance of this peptide for microvascular repair in ALI and ARDS has not been explored.

It has increasingly been recognized that ECs play a key role in orchestrating tissue repair through the self-renewal and differentiation of resident stem and progenitor cells in an organ-specific manner(16). Recently the understanding of this heterogeneous EC landscape has been greatly facilitated by the introduction of single cell transcriptomics analysis. Using this approach, two specialized lung microvascular EC populations have recently been described in the normal lung(17); ‘aerocytes’ (aCap ECs), which are characterized by the expression of apelin, and general (gCap) ECs expressing the apelin receptor. Aerocytes are highly differentiated, large cells which make up the endothelial component of the alveolar air blood barrier and are incapable of proliferation(17). In contrast, gCap ECs are smaller and located at thicker regions of the alveolar wall(17) and respond to injury by proliferation; therefore, represent the lung EC population within which endothelial stem cells may arise.

Using a new model of ALI induced by targeted lung EC ablation, we now demonstrate that EC regeneration is initiated by the emergence of a new gCap endothelial stem cell population post injury that paradoxically exhibits the *de novo* expression of apelin together with a stem cell marker, protein C receptor (*Procr*). This population rapidly transitioned to highly proliferative progenitor-like cells, characterized by apelin receptor and *FoxM1* expression, which were responsible for repopulating all depleted lung endothelial fields, including aerocytes, leading to rapid ALI resolution by an apelin-dependent mechanism.

## Methods

### Animal model

Transgenic Cre-inducible DT animals were obtained from Jackson’s Laboratories R and only male binary transgenic animals 10-12 weeks of age were used for experiments (Figure 1A) in this manuscript, unless otherwise specified. All animals were genotyped using primer sequences provided by Jackson’s Laboratory. Animals were anaesthetized and DT was delivered by intra-tracheal (IT) instillation (10 ng in 50 μLs, unless otherwise specified). Control animals received 50 μLs of 0.9% sterile saline.

**Figure 1.**
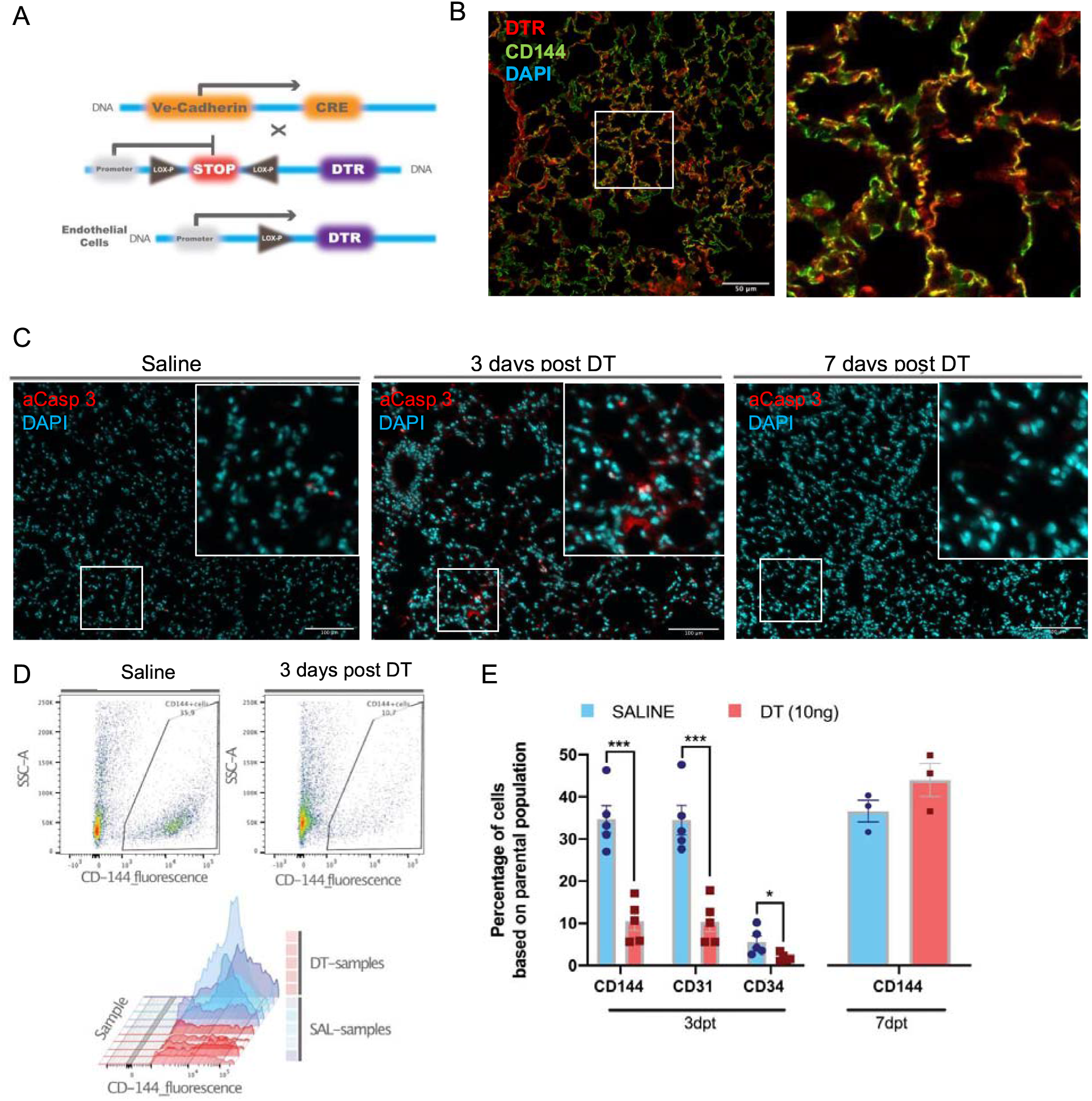
Establishment of the DT-induced EC ablation model: A) Transgenic mice harboring Cre recombinase cDNA downstream from a 2.5-kb fragment of the VE-Cadherin (Cdh5) mouse promoter (B6.FVB-Tg(Cdhn5-cre)7Milia) were crossed with mice with Cre-inducible expression of DTR (C57BL/6-Gt(ROSA)26Sor^tm1(HBEGF)Awai^/J) giving rise to Cdhn5-DTR binary transgenic mice. B) Immunostaining for human DTR (red) was associated with ECs (anti-CD144, green) in lung sections from Cdhn5-DTR binary transgenic mice 3 days post IT delivery of saline or DT (10ng) with DAPI nuclear staining (blue). C) Immunostaining for activated caspase 3 (aCasp-3, red) in lung sections from binary transgenic mice 3 or 7 days after treatment DT or saline (DAPI nuclear staining in blue). D) Representative plots of lung EC numbers assessed by flow cytometry (CD144) in binary transgenic mice 3 days after IT delivery of saline or DT. E) Summary flow cytometric data showing the percent of total lung cells staining positive using CD144, CD31 or CD34 antibodies in DT treated binary transgenic mice 3 or 7 days post treatment (dpt) compared with saline. * = p<0.05; ** = p<0.01; *** = p<0.005

### Single-Cell RNA Sequencing Analysis

Details are provided in the online supplement. In brief, RNA library was constructed with 10x-Genomics Single Cell 3’ RNA sequencing kit v3 as previously described(18) sequenced using a NextSeq500 (Illumina). Cell clusters were characterized using an automated annotation tool(19) and by cross referencing differential gene expression of individual clusters to previously characterized lung cells of the *Tabula Muris* cell atlas(20). The identity of the various lung cell clusters was further confirmed by the assessment of expression of cell-specific genes in the 21 clusters (Figure S5 and S6).

### Apelin inhibition

The role of apelin pathway in lung microvascular repair was assessed using a selective apelin receptor antagonist, ML221 (Tocris, 10mg/kg)(21) dissolved in DMSO delivered by intraperitoneal injection as previously described(22).

#### Statistical analysis

Statistical analysis was performed using Prism v.8.4.2 (GraphPad Software). Results are expressed as means ± SEM. The sample size per condition and specific statistical tests used are presented in the figure legends. Statistical analysis for the scRNA-seq data was performed according to the recommendations of analytic packages that were used as specified in the Results section and figure legends.

## Results

### Selective lung EC ablation in Cdh5-DTR mice

Administration of DT at doses below 20 ng IT was consistent with survival (Figure S1A) and resulted in modest increases in right ventricular systolic pressures (RVSP) that were maximal at 10ng of DT (Figure S1B). DTR expression was localized to ECs in Cdh5-DTR animals (Figure 1B) and widespread EC apoptosis was seen at 3 days post DT administration (10ng) in Cdh5-DTR mice (Figure 1C). Indeed, DT administration resulted in a ~70% reduction in EC numbers at 3 days by flow cytometry (p<0.001) (Figure 1D and E) with full recovery by day 7 (Figure 1E). Lung permeability was increased (p<0.01) at 3 days post DT administration (Figure 2A and B), again returning to normal by 7 days, with no evidence of increased vascular permeability in other organs (Figure 2B and Figure S2). At 3 days there was a marked increase in the lung injury score (Figure 2C and D) consistent with severe ALI, and this was associated with the influx of CD11b+Ly6G+ neutrophils by flow cytometry (Figure 2E).

**Figure 2.**
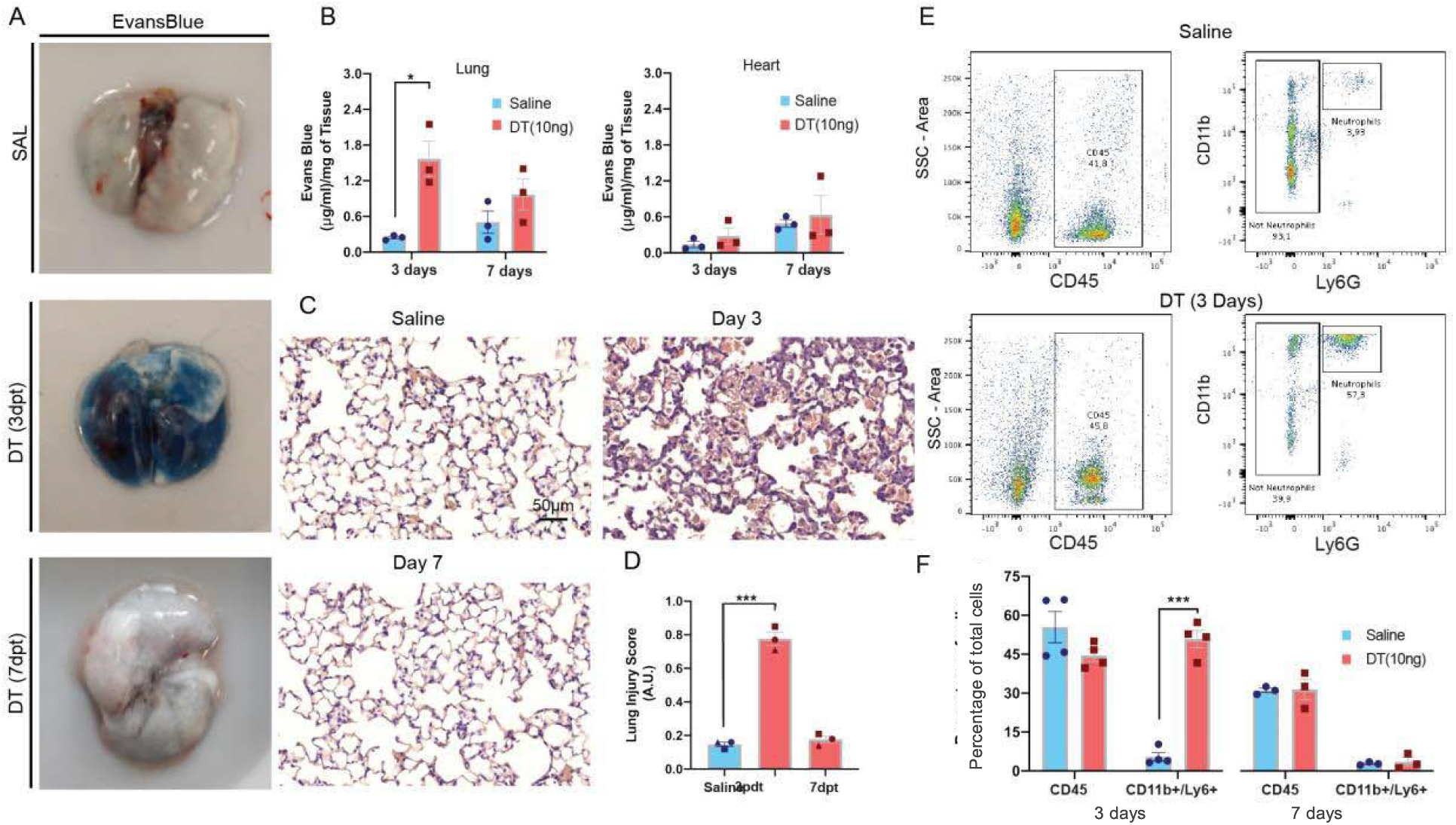
DT administration results in an acute lung injury phenotype. A) Representative examples of Evans blue staining of lungs from binary transgenic mice. B) Summary data showing marked increase Evans Blue lung content in binary transgenic mice treated with DT (10ng) or saline at 3 and 7 days post treatment (dpt). C) Representative histological lung sections (H&E staining) from binary transgenic mice at 3 and 7 days post IT administration of DT or saline with the corresponding summary data (D) for a validated lung injury score. E) Representative examples of flow cytometry plots showing gating strategy for assessing CD11b/Ly6G positive leukocytes in lungs from binary transgenic mice 3 days after IT delivery of DT or saline with summary data (F) at 3 and 7 days post treatment (dpt). An unpaired multiple t-test with Holm-Sidak multiple comparisons method with alpha (0.05) was used for analysis of data presented in panels B, D and F, whereas an unpaired t-test was conducted for panels C-E. * = p<0.05; ** = p<0.01; *** = p<0.005

### Changes in global lung cell populations with single cell transcriptomic profiling

Multiplexed scRNA-seq analysis was performed on lung tissues of Cdh5-DTR mice at baseline (day 0) and 3, 5 and 7 days post DT administration (Figure 3A). Cells lacking a barcode, or positive for multiple barcodes were excluded from further analysis (Figure S4A). Uniform Manifold Approximation and Projection (UMAP) dimensionality reduction maps of the data revealed 21 separate cell populations, representing immune, endothelial, stromal and epithelial cells (Figure 3B). While EC populations, together with type 2 alveolar (AT2) epithelial cells and alveolar macrophages, were reduced after DT administration, there was an increase in inflammatory cells, notably neutrophils and monocytes at days 3 and 5, respectively, all recovering by day 7 (Figure 3C). Cell types that were most affected by DT treatment were identified using Augur(23) and included ECs (clusters 11 and 0), classical monocytes (cluster 4), alveolar macrophages (cluster 5) and type 2 pneumocytes (cluster 10) (Figure 3D).

**Figure 3.**
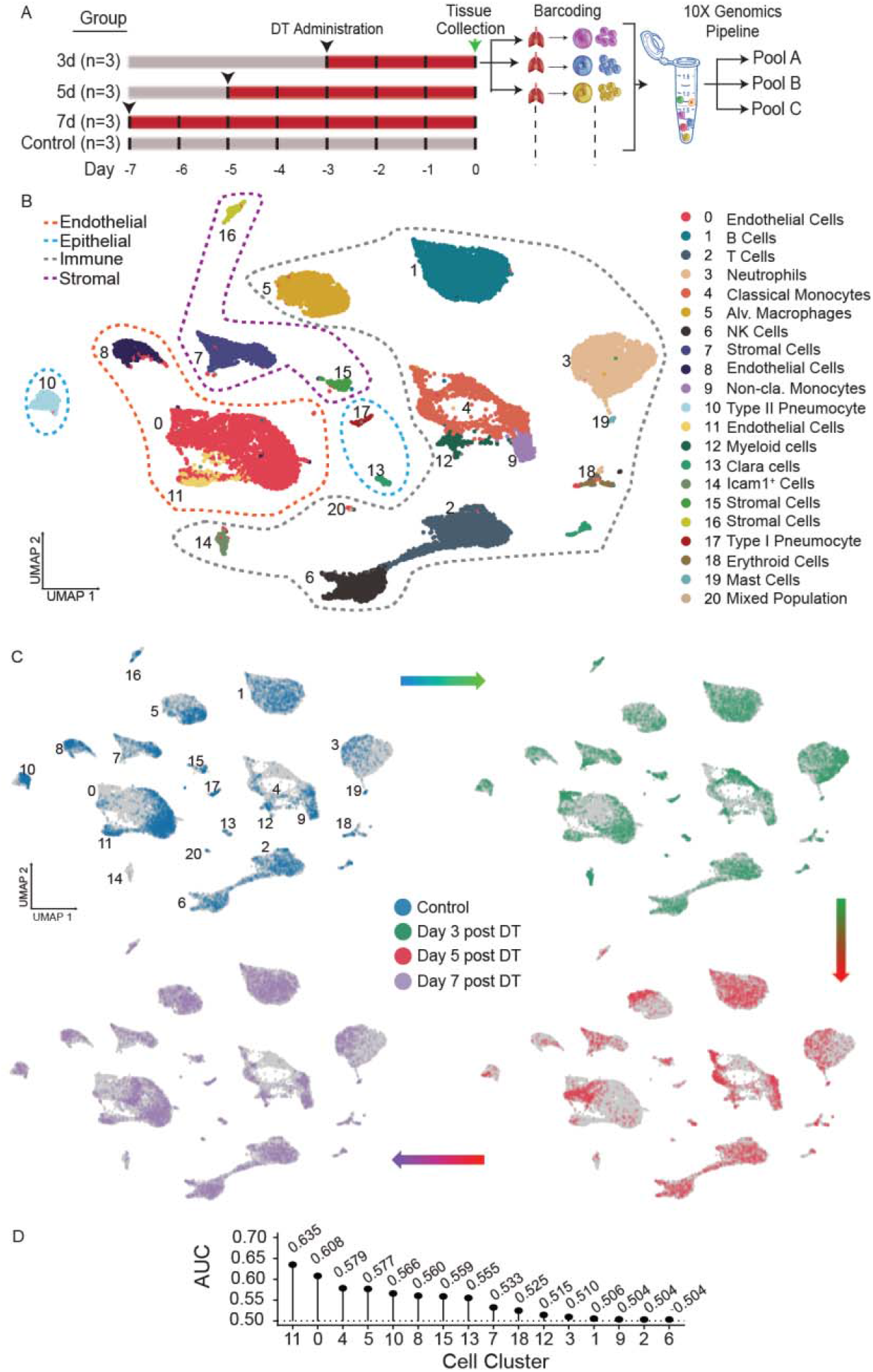
Multiplexed single cell transcriptomic analysis after DT-induced lung EC ablation: A) Schematic of workflow showing the experimental design. Three separate cohorts (3, 5 and 7 days) of binary transgenic mice received DT (IT, 10ng), and a fourth cohort of healthy animals served as a control, with timing of DT delivery such that all mice were sacrificed on the same day. Lungs cells were immediately isolated and barcoded to identify individual donor animals, then pooled and subjected to library construction using 10x-Genomics Single Cell 3’ RNA sequencing kit v.3. Global plot of all lung cells at all time points using uniform manifold approximation and projection (UMAP). B) 21 distinct populations were identified and could be assigned in to four major categories: immune; endothelial, stromal and epithelial. C) UMAP plots of global lung cells are shown for each mouse cohort: control (blue); day 3 (green) day 5 (red) and day 7 (purple) (C). D) A machine learning model was used to predict cells that become more separable during treatment based on their molecular measurements (https://github.com/neurorestore/Augur).

### Changes in EC populations in response to injury

Re-clustering of the EC populations yielded 8 distinct clusters (Figure 4A), including five capillary populations (i.e. clusters 0, 1, 3, 6 and 7) which exhibited gene expression profiles typical for alveolar gCap ECs (Figure S7),(17,24) and cluster 2 that corresponded to aCap ECs also called aerocytes. Pulmonary arterial and venous ECs (clusters 4 and 5, respectfully) also expressed typical gCap genes, consistent with the general nature of this expression profile. Cluster 1 was unique in that it was made up of 4 distinct ‘zones’; each zone specific to a single time point before and after EC ablation (Figure 4B), whereas cluster 7 emerged mainly at 5 days after EC injury. Most EC populations, notably aCap ECs, exhibited a marked decline in cell numbers after DT-induced EC ablation, reaching a nadir of almost 90% cell loss at 5 days (Figure 4C and D). Only clusters 1 and 7 showed stable or even increased cell numbers, consistent with a role of these EC populations in lung microvascular repair.

**Figure 4.**
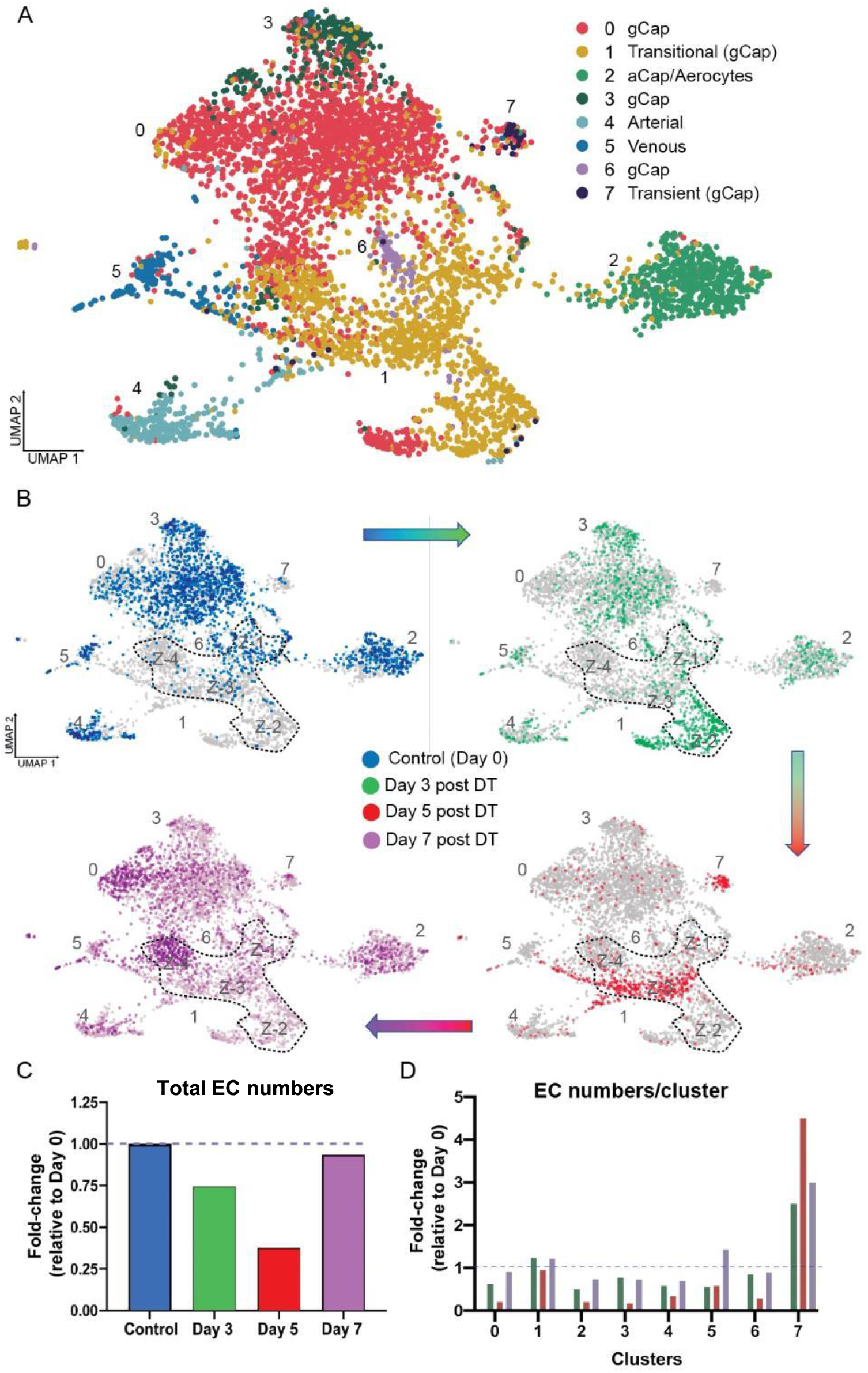
Endothelial cell populations in control and DT treated binary transgenic DTR mice: A) UMAP representation of the re-clustered endothelial populations for all cohorts produced 8 distinct EC clusters. B) The UMAP plots for each cohort are shown separately: control (blue); day 3 (green) day 5 (red) and day 7 (purple) The dotted line delineates cluster 1 which was unique in that it consisted of four zones (Z-1 to Z-4) each specific to a single time cohort. C) The total numbers of lung ECs in the three cohorts treated with DT (Days 3, 5 and 7) are expressed as fold-change relative to control for all EC populations. D) Numbers of ECs broken down for each EC cluster.

### Temporal sequence of angiogenic gene expression in aCap and gCap ECs

At baseline (day 0), apelin was found to be uniquely expressed in aCap ECs of cluster 2 (Figure 5A) as previously reported,(17,25) whereas apelin receptor (*Aplnr*) was found only in gCap ECs (Figure 5A). In addition to apelin, aCap ECs also exhibited predominant expression of other EC tip cell genes including *Kdr (Vegfr2), Npl1* (neuropilin 1) and *Cd34* (Figure S8A),(26,27) whereas stalk cell markers, such as *Hey1* (Figure S7B), *Flit1 and Notch1* (Figure S8B) were mainly expressed by gCap ECs. Three days after EC injury, *de novo* expression of apelin was seen in the newly emergent gCap EC population in zone 2 of cluster 1 (Figure 5A). Despite the expression of all other gCap markers, these cells lacked *Aplnr* and exhibited the expression of *Procr* (protein C receptor or EPCR), a marker of bipotent resident vascular endothelial stem cells(28). Using two gene transcriptomic analysis, co-expression of apelin with *Procr*, as well as *Cd93* was evident at 3 days post injury as confirmed by immunofluorescence staining (Figure 5C). To our knowledge, this represents the first demonstration of apelin expression by gCap ECs. As well, *Angpt2* (angiopoietin 2) was uniquely expressed in this endothelial stem cell-like population,(29) together with the progenitor cell marker, *Cd34* (Figure S8A). At day 5 post DT, new populations emerged in zone 3 of cluster 1, as well as in cluster 7 (Figure 4B) exhibiting *Aplnr* expression together with *Ki67*, a marker of cell proliferation (Figure 5A), and *FoxM1*, a pro-proliferative transcription factor (Figure S8B); however, the density of proliferating cells was greatest in cluster 7. By day 7, co-expression of apelin and CD93 by single cell transcriptomics and immunostaining was rarely seen and the expression of *Procr*/EPCR was nearly entirely lost.

**Figure 5.**
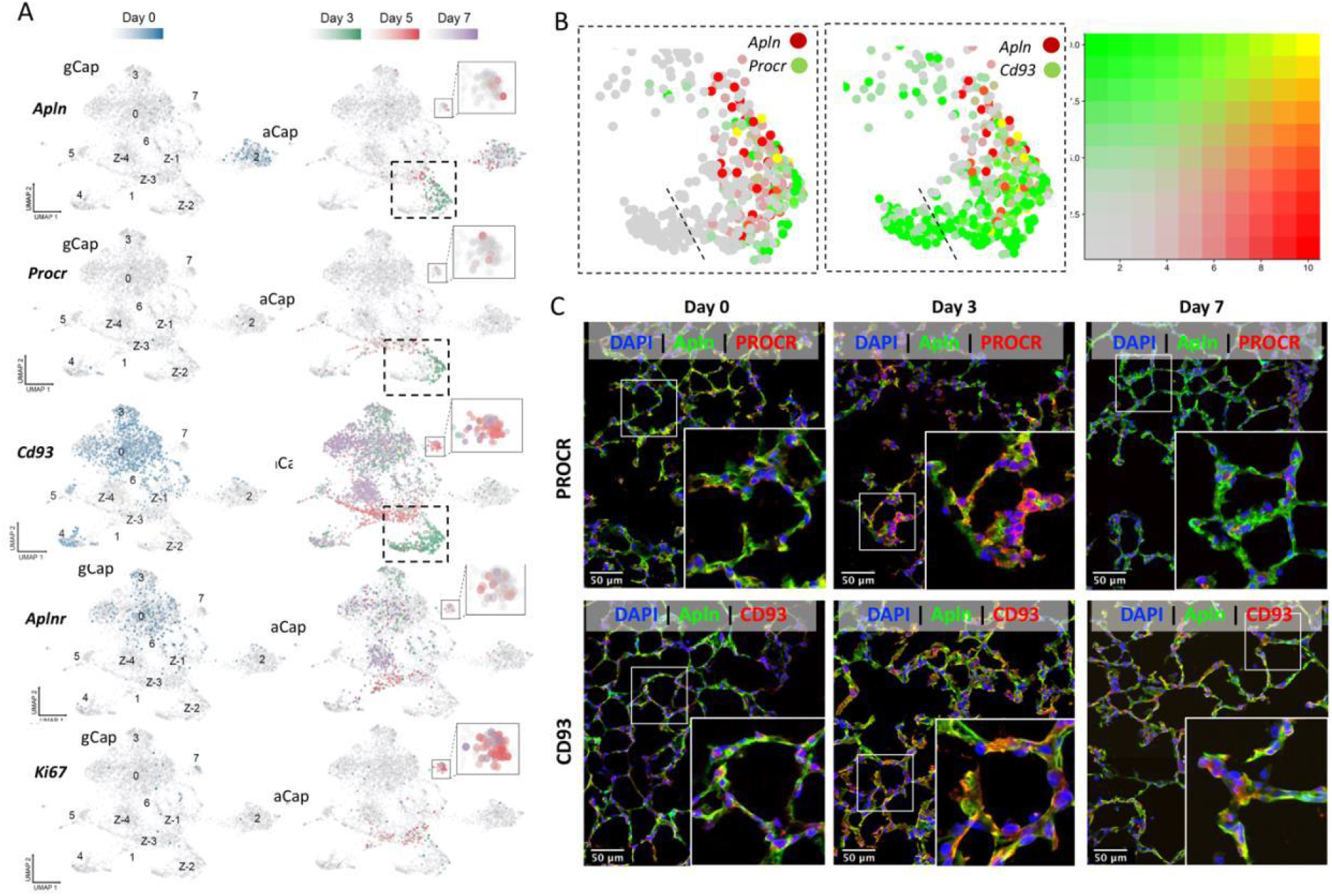
Apelin expression in a transitional gCap stem-like EC population at 3 days post injury: A) Under basal conditions (Day 0; blue color) apelin was only expressed by aCap ECs (aerocytes; cluster 2), whereas *Aplnr* expression was seen in gCap ECs (*Cd93* positive). 3 days after EC ablation (Green color), there was de novo apelin expression in gCap ECs of zone 2, cluster 1, together with *Procr*; while *Aplnr* and *Ki67* in the adjacent zone 3 of cluster 1 and in cluster 7 at 5 days post EC injury (Red colour). B) Enlargement of cluster 1 showing expression of Apelin, *Procr* and *Cd93* in zone 2 at 3 days. C) Immunofluorescence staining showing colocalization of APLN, PROCR and CD93 in zone 2 of cluster 1only at 3 days post EC ablation.

### Differential gene expression and pathway enrichment analysis of endothelial clusters

Figure 6A shows volcano plots of differentially expressed genes (DEGs) compared to cluster 0 and the top 10 DEGs are presented in Figure 6B. For this analysis, only zone 1 of cluster 1, which represents Day 0, was included. Not surprisingly, this cluster exhibited a high degree of similarity in gene expression profile with cluster 0, the largest gCap population, although there was an increase by pathway enrichment analysis in activation of pathways associated with oxidative phosphorylation, as well as EIF2 signaling which was also shared by clusters 4 and 5 (arterial and venous ECs, respectively) (Figure 6C). As expected, cluster 2 showed high expression of typical aCap genes, such as *Endrb, Fibin and Car4*, and pathways related to PKA and Ephrin B signaling were enriched. Interestingly, cluster 3 showed upregulation in many early response genes, including *Erg1, Junb, Ier2, Ier3* and *Fosb* with enrichment in signaling associated with a number of inflammatory pathways; whereas cluster 6 exhibited no predicted pathways with significant activation or inhibition. EC cluster 7, showed the greatest number of DEGs (~1200), nearly all of which were upregulated, including many genes involved in cell proliferation such as *Rrm2, Aurkb, Cdc25c, Tk1, Cdca8* and *Birc5*, and a unique activation of signaling pathways involved in cell cycle regulation. The identities of all significantly enriched canonical pathways are provided in supplementary Figure S9.

**Figure 6.**
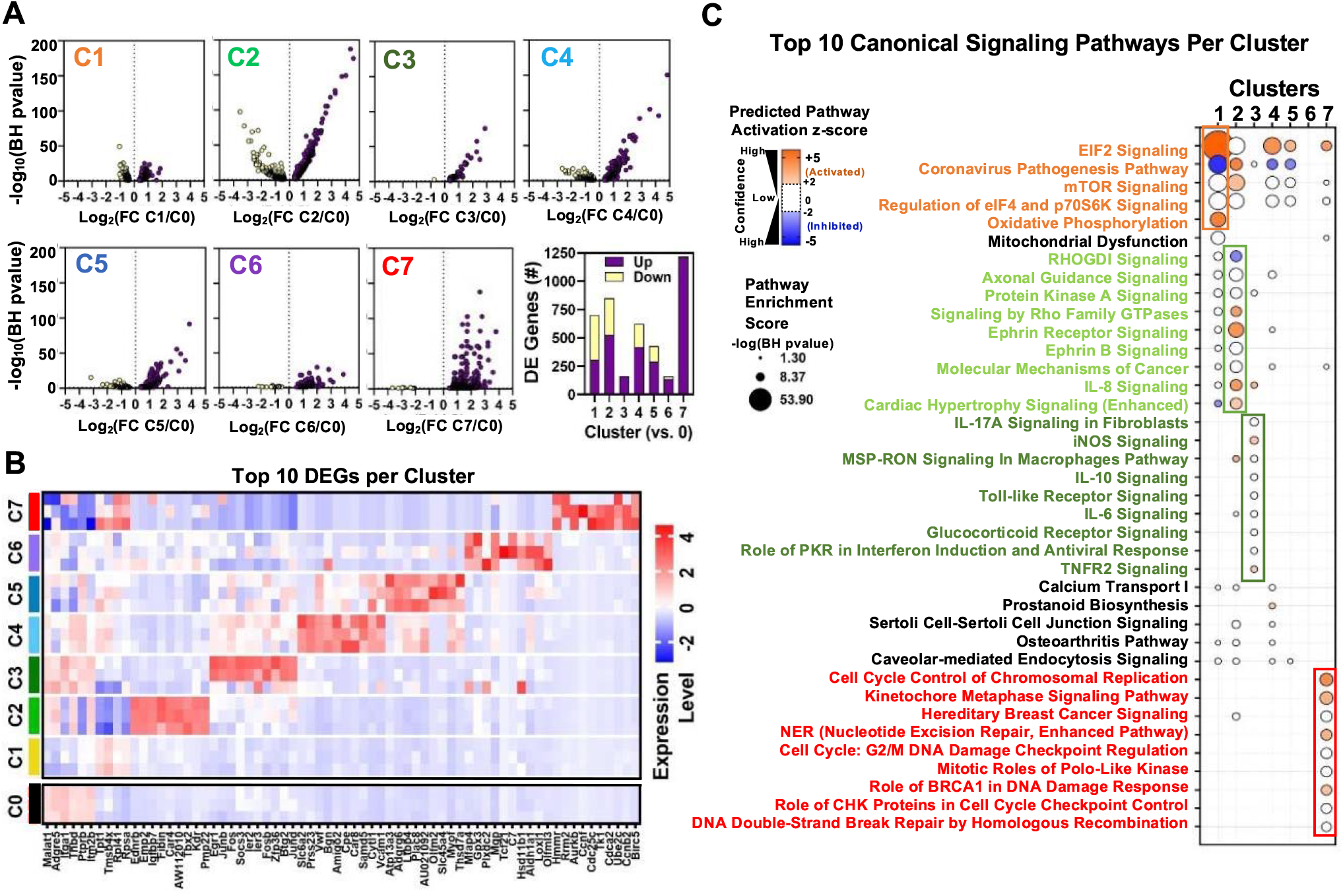
Differential gene expression and pathway enrichment analysis of endothelial sub-clusters 1-7. **A)** Volcano plots showing distribution of differentially expressed genes in clusters 1-7 (C1 to C7) according to statistical significance (Benjamini-Hochberg (BH) adjusted p-value), magnitude and direction of fold change (FC) relative to cluster 0 (at baseline time 0). Only genes that passed the false discovery rate threshold (FDR) <0.05 are shown. Bar chart summarizes of the number of differentially expressed (DE) genes in each cluster (versus cluster 0). Purple and yellow bars are stacked to show the proportion of genes that were upregulated and downregulated, respectively. B) Heatmap shows expression levels of 68 genes representing the combined top 10 differentially expressed genes in clusters 1 to 7 (versus cluster 0) at baseline day 0. Mean expression levels (z-score of log transformed normalized counts) are shown for n=3 mice/cluster. C) Pathway enrichment analysis using DE gene sets from panel A with Ingenuity Pathway Analysis. Identity of 38 canonical signaling pathways representing the combined top 10 most significantly enriched pathways in each endothelial subcluster and the associated overlap between clusters. Of note, some pathways may be classified in more than one cluster. Circle size denotes pathway enrichment score based on the BH adjusted p-value. A minimum score of 1.3 (i.e., FDR<0.05) was used as an inclusion threshold. Z-score color denotes the predicted activation state of the pathway based on the degree of matching between the expected and observed pattern of gene expression changes. Only pathways with z-score >+2 (activated) or <-2 (inhibited) are shown in color to highlight confident predictions. White circles denote pathways in which the activation state cannot be confidently predicted. No pathways were significantly enriched in DE genes from Cluster 6 at FDR<0.05. The identities of all significantly enriched canonical pathways are presented in **Supplementary Figure S9.**

The fact that cluster 1 was comprised of four distinct zones, each specific to a single timepoint, afforded a unique opportunity to assess the differential gene expression in response to EC ablation within a single cluster spanning the three distinct phases of microvascular regeneration and repair. Zones 2 and 3 (days 3 and 5) both showed high numbers of DEGs (~800-1000) compared with zone 1 (Day 0) (Figure 7A and B), and many of these were unique to these zones (Figure 7C). In contrast, there were few DEGs in zone 4, consistent with relative normalization of gene expression profiles by 7 days post injury. A heatmap showing the top 20 DEGs relative to zone 1 (Day 0) revealed upregulation of genes related to cell growth and response to injury in zone 2 (Figure 7D), some of which were unique to this zone (*Odc1* and *Plat*), while others were shared with zone 3 (*Ccdn1* and *RhoC*). Zone 3 also exhibited unique upregulation of genes involved in cell proliferation and survival (*Birc5*, *Malat1* and *Jpt1*), as well as metabolism (*Pkm*); whereas DEGs in zone 4 were mainly related to differentiation (*Sox4, Sparc1* and *Hlx*) and cell cycle inhibition (*Rgcc*). The top 10 signaling pathways by pathway enrichment analysis showed increases EIF2, eIF4 and integrin signaling in zone 2 ECs as well as pulmonary and hepatic fibrosis signalling (Figure 7E). There was further enrichment of EIF2 and mTOR signaling in Zone 3 along with unique upregulation in oxidative phosphorylation. In contrast, there was no significant predicted activation of signalling pathways in zone 4 (Day 7) compared with baseline (zone 1). The identities of all significantly enriched canonical pathways are presented in supplementary Figure S10.

**Figure 7.**
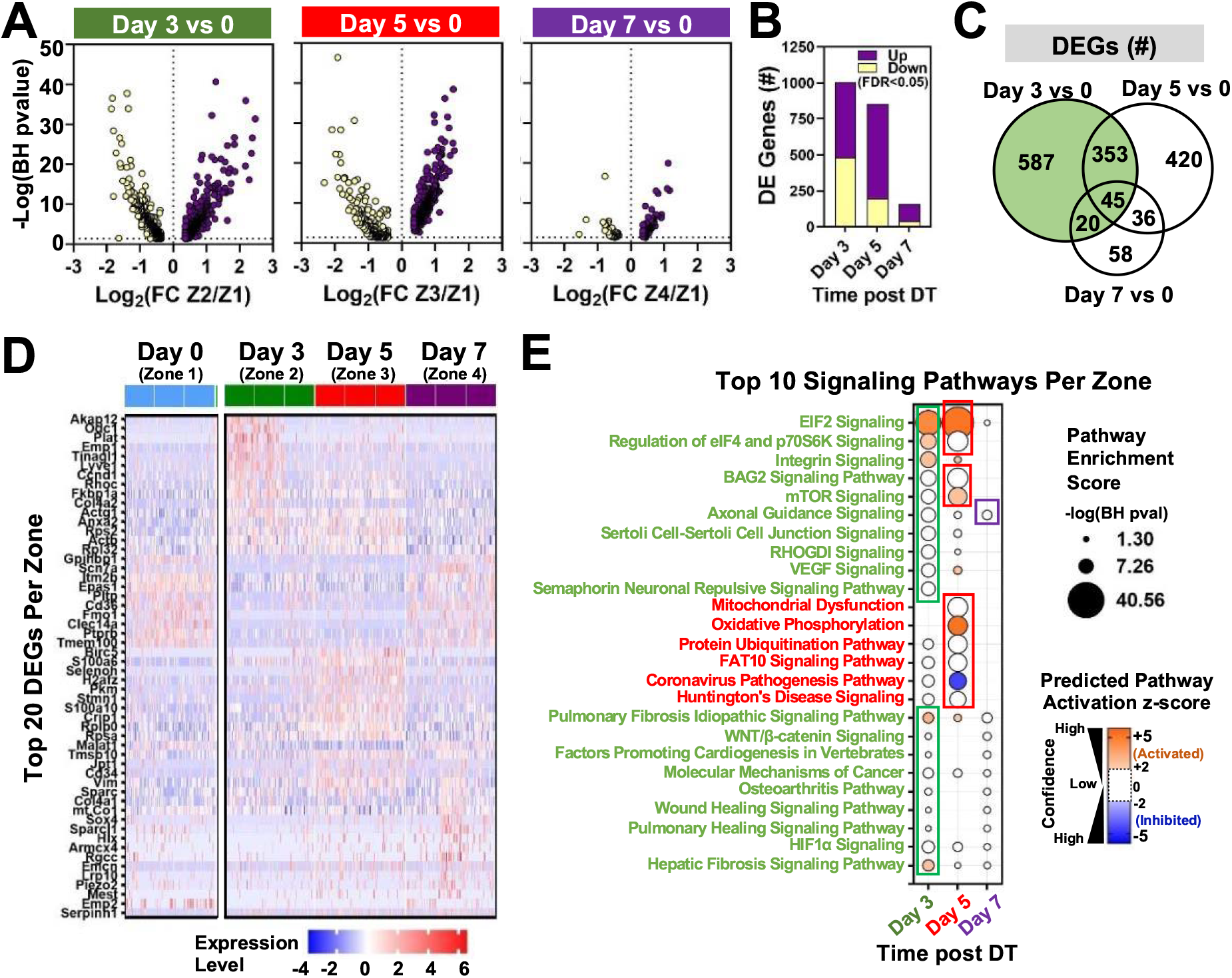
Differential expression and pathway enrichment analysis of time-dependent changes in zones 1-4 of endothelial cluster 1. **A)** Volcano plots showing distribution of differentially expressed genes according to statistical significance, magnitude and direction of fold change (FC) at different points following administration of diphtheria toxin (DT) in mice. Only genes that passed the false discovery rate (FDR) threshold <0.05 are shown (i.e., Enrichment score -log(BH pvalue)>1.3). Zones 1, 2, 3, and 4 denote day 0, day 3, day 5, and day 7, respectively. B) Summary of the number of differentially expressed (DE) genes in each comparison. Purple and yellow bars are stacked to show the proportion of genes that were upregulated and downregulated, respectively. C) Venn diagram shows degree of overlap in the identity of differentially expressed genes for each comparison. D) Heatmap shows expression levels of 54 genes representing the top 20 differentially expressed genes at days 3, 5 and 7 versus day 0. Expression levels (z-score of log transformed normalized counts) are shown for a sample of 50 cells from each mouse (n=3 mice/time point). E) Pathway enrichment analysis using DE gene sets from panel B with Ingenuity Pathway Analysis. The top 10 signaling pathways (by enrichment score) are shown for each time zone. Circle size denotes pathway enrichment score based on the Benjamini-Hochberg (BH) corrected p-value. A minimum score of 1.3 (i.e., FDR<0.05) was used as an inclusion threshold. Z-score color denotes the predicted activation state of the pathway based on the degree of matching between the expected and observed pattern of gene expression changes. Only pathways with z-score >+2 (activated) or <-2 (inhibited) are shown in color to highlight confident predictions. White circles denote pathways in which the activation state cannot be confidently predicted. The identities of all significantly enriched canonical pathways in zones 1-4 of cluster 1 are presented in **Supplementary Figure S10**.

### ALI resolution and survival is dependent on apelin signalling

An analysis of a publicly available atlas of the aging lung(30) revealed a significant reduction in lung apelin expression in mice (Figure 8A) which was confirmed in the Cdh5-DTR double transgenic mice by RT-qPCR (Figure 8B). Interestingly, aging reduced survival post DT-induced injury with only 24% of 52 week-old Cdh5-DTR mice surviving to day 7 post DT compared to 100% survival for 12 week-old mice (p<0.01) (Figure 10C). Moreover, when young Cdh5-DTR mice were treated with an apelin receptor antagonist, ML221, all mice succumbed to DT-induced acute lung injury by 5 days (Figure 8C), consistent with a failure of lung microvascular repair.

**Figure 8.**
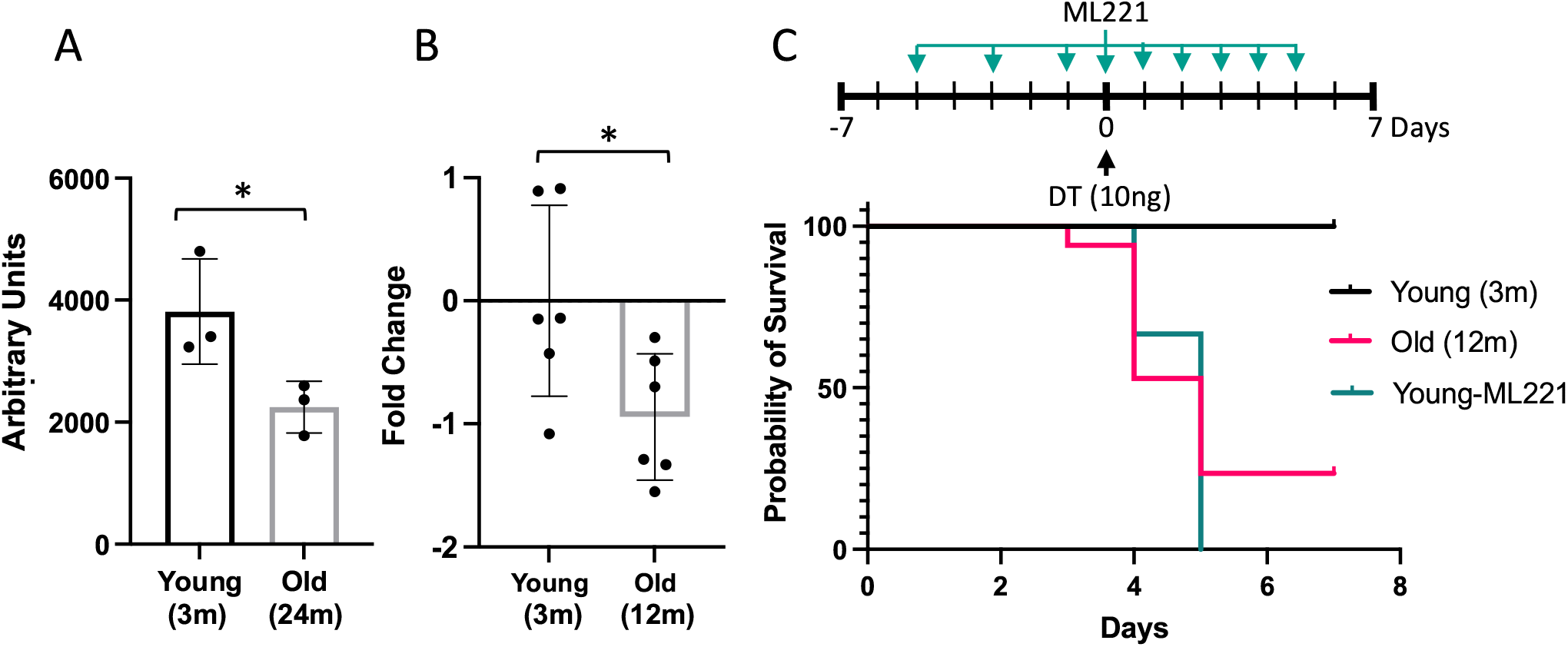
Impairment or inhibition of the apelin pathway leads to failure of recovery after EC ablation: **A**) Expression of apelin from a publicly available dataset (31) in young (3 months) versus old (24 months) mice (* = p<0.05, unpaired t-test). **B**) Apelin expression assessed by qRT-PCR expressed as fold-change relative to mean value in young (3 month) Cdhn5-DTR binary transgenic mice (* = p<0.05, unpaired t-test). **C**) Survival after IT instillation of DT of either 3 (Young) or 12 month-old (Old) Cdhn5-DTR with or without administration of the apelin antagonist, ML221 (10mg/kg IP). Survival of old mice or mice receiving ML221 was significantly different from young control mice (p<0.05, Mantel Cox log rank test).

## Discussion

In this study, we have established that selective ablation of lung ECs results in a severe ALI model with many features of ARDS seen in various clinic contexts, including COVID-19. Remarkably, the loss of more than 70% of lung ECs was compatible with survival by virtue of an efficient endogenous regenerative response that resulted in near complete restoration of the lung microvascular structure and function in just one week. This model underscores the importance of the endothelium in ALI and affords a unique opportunity to explore the mechanisms underlying EC regeneration and microvascular repair, which is critical for ALI resolution. Using single cell transcriptomic analysis, we have identified novel regenerative EC populations emerging in a tightly scripted temporal sequence after EC injury. We now show for the first time the robust expression of apelin within gCap, stem-like ECs that give rise to apelin receptor positive, highly proliferative progenitors which are responsible for the replenishment of all depleted EC pools, including the highly specialized aerocytes (aCap ECs) that reform the air-blood barrier.

In the normal uninjured lung, we confirmed that the expression of apelin identified a EC population corresponding to alveolar aCap ECs, or ‘aerocytes’, as previously described by Gillich et al.(17). These were characterized as large, highly specialized cells forming the air-blood barrier, thereby playing an important structural role in gas exchange. This population also appears to be analogous to an EC population described earlier in close apposition to alveolar type 1 epithelial cells and expressing high levels of Car4,(24,25) one of five genes typically expressed by aCap ECs.(17,25) We also describe, for the first time, the emergence of novel EC populations after EC injury which, based on expression of typical marker genes, originate from gCap ECs. In particular, we have identified a gCap EC population that, remarkably, exhibits *de novo* expression of apelin together with gCap markers such as CD93 (with the exception of the apelin receptor) demonstrating that apelin expression in the alveolar capillary bed is not necessarily restricted to aCap ECs. This population was also characterized by the unique expression of two other genes, *Procr* (protein C receptor) and *Angpt2* (angiopoietin 2), that have both been implicated in angiogenesis and vascular repair. Angiopoietin 2 is an endogenous antagonist of the Tie2 receptor that is strongly expressed in endothelial tip cells(31) and is instrumental in the initiation of angiogenic response(29), whereas *Procr* has recently been reported to be a marker of bipotent resident vascular endothelial stem cells that have the capacity to regenerate ECs as well as pericytes(28), the two cell types required for the genesis of stable neovessels. Interestingly, expression of *Procr* also identifies a subpopulation of CD34^+^ haematopoietic stem cells (HSCs) with markedly greater proliferative and bone marrow engraftment potential(32). Moreover, *Procr* has been suggested to be a marker of primitive, endothelial-like hematopoietic precursors (i.e. pre-HSCs)(33), that give rise to both ECs and HSCs during development(34); therefore, playing a critical role in the early development of both blood and blood vessels. Together, this is consistent with the stem cell nature of this transient population of endothelial stem cells which initiates the regenerative response after EC ablation.

Apelin-positive, gCap ECs showed no evidence of proliferation at 3 days post injury; however, by 5 days they transitioned to a highly proliferative phenotype with expression of apelin being replaced by its receptor, consistent with a dynamic role for the apelinergic system in driving a transition from endothelial tip-to stalk-like, proliferative progenitor cells(35). At the same time, a very similar stalk-like progenitor population emerged within the transient EC cluster 7. Both these populations expressed *FoxM1*, which has previously been implicated in lung microvascular repair(9), and exhibited similar proliferative gene expression signatures; however, cluster 7 showed a greater density of *FoxM1* and *MKi67* expressing proliferative ECs. Finally, the pivotal role of apelin in EC regeneration was confirmed by the failure of resolution of ALI after DT-induced lung injury and excessive mortality in older mice exhibiting reduced expression of apelin and with the administration of a selective receptor antagonist.

Thus, we have delineated the cellular and molecular processes involved in rapid and complete microvascular regeneration using a new model of ALI induced by targeted EC ablation. Single-cell transcriptomic analysis has revealed a novel gCap endothelial stem-like population expressing apelin and *Procr* and emerging soon after endothelial injury, subsequently giving rise to *FoxM1*-positive progenitor-like ECs expressing the apelin receptor. These highly proliferative ECs are responsible for replenishing all depleted EC populations leading to the rapid resolution of the ALI phenotype by an apelin-dependent mechanism. These findings highlight a critical role of apelin signaling in regulating the activity of regenerative cells that mediate lung microvascular repair and provide insights for the development of novel regenerative strategies for the treatment of ALI and ARDS.

## Acknowledgments

Would like to thank Dr. Saad Khan, Dr. Maria Hurskainen and Dr. Ivana Mizikova for procedural guidance and advice and to Dr. Bernard Thébaud for his advice on the manuscript.

## Author contributions

RSG has contributed to all aspects of this work, from conception and performance of the experiments to data analysis and drafting of the manuscript. DPC was instrumental in the performance, analysis, and interpretation of the single-cell RNA-seq studies. NDC contributed to lung dissociation optimization, single cell study acquisition, flow cytometry acquisition, the performance and interpretation of in vivo studies, and manuscript revisions. EM contributed to analysis and interpretation of single-cell RNA-seq studies. YD was involved with animal studies and with the performance and analysis of the micro-CT data. LW performed immunostaining and contributed to the analysis of the single-cell transcriptomics data and drafting of the manuscript. AC contributed to studies on inhibition of apelin signaling. KS performed differential gene expression analysis and gene ontology analysis, data interpretation, and manuscript preparation.

KR was involved with animal studies, flow cytometry panel design, flow cytometry acquisition, data analysis and interpretation. DJS was involved in conception and design of experiments, data analysis and interpretation and drafting of the manuscript, and is accountable for all aspect of this work.

## Non-standard Abbreviations and Acronyms

ALI: Acute lung injury
ARDS: Adult Respiratory Distress Syndrome
aCap: Aerocytes
DGE: Differential gene expression
DT: Diptheria toxin
DTR: Diptheria toxin receptor
ECs: Endothelial cells
FoxM1: Forkhead box M1
gCap: General capillary
IT: Intra-trachael
RV: Right ventricle
RVSP: Right ventricular systolic pressure
scRNA-seq: Single-cell RNA sequencing
UMAP: Uniform manifold approximation and projection

