## Supplemental Data for "Novel Apelin-expressing gCap Endothelial Stem-like Cells Orchestrate Lung Microvascular Repair"

**Online Supplement**

**Supplemental Methods**

**Transgenic models**: All animal procedures were approved by the University of Ottawa Animal Care Ethics Committee in agreement with guidelines from the Canadian Council for the Care of Laboratory Animals. Transgenic animals were obtained from Jackson's Laboratories including mice with Cre-inducible DTR flanked by loxP sites (C57BL/6-*Gt(ROSA)26Sor*$^{tm1(HBEGF)Awai}$/J) and expressing Cre-recombinase under the EC restricted promoter, VE-cadherin (Cdh5) (B6.FVB-Tg(Cdh5-cre)7Mlia/J) (Stock numbers: 007900 and 006137, respectively). Stocks were maintained by crossing homozygous animals. Binary transgenic animals harboring Cdh5-cre-iDTR were obtained by crossing homozygous DTR and Cdh5 mice to generate double heterozygous offspring (Figure 1A). Only male animals 10-12 weeks of age were used for experiments in this manuscript, unless otherwise specified. All animals were genotyped using primer sequences provided by Jackson's Laboratory.

**Diphtheria Toxin Administration**: Stocks of Diphtheria toxin from *Corynebacterium diphtheriae* (Sigma-Aldrich, Oakville, ON, Canada) were prepared (1 mg/mL) in sterile distilled water and stored in single use aliquots at -20 ºC. Animals were anaesthetized with ketamine (100 mg/Kg i.p.) and xylazine (10 mg/Kg i.p.) and DT was delivered by intra-tracheal (IT) instillation (10 ng in 50 µLs, unless otherwise specified) with a 1 mL syringe and measured with a micropipette. Prior to toxin administration, a water droplet was used to indicate proper positioning of the catheter into the orotracheal cavity and deemed successful by the movement of the droplet with the breathing of animals. Control animals received 50 µLs of 0.9% sterile saline.

**Lung dissection and single cell dissociation**: Animals were anaesthetized with ketamine (100 mg/kg) and xylazine (10 mg/kg) (i.p.) and given 10 mU/g heparin sodium (i.p.) (LEO Pharma Inc., Thornhill, ON, Canada). The inferior vena cava was exposed surgically and the animals were bled and then 25 U/mL heparin in 10 mLs of sterile 0.9% saline was flushed through lungs via pulmonary artery until they were cleared of all evidence of red blood cells. Lungs were rinsed in PBS, cut into smaller pieces and placed into gentleMACS C tubes™ (Miltenyi Biotech, Bergisch Gladbach, Germany) containing 2.5 mLs DPBS to which 2.5 mLs of digestion enzyme mix were added. This mix included 2500 U Collagenase I, 30 U Neutral Protease/Dispase (Wothington Biochem., Lakewood, NJ, USA), and 500 U Deoxyribonuclease (Sigma-Aldrich, Oakville, ON, Canada) in 1x Dulbecco's PBS (Thermofischer Scientific, Burlington, ON, Canada), and was made fresh for each experiment and kept on ice. GentleMACS C tubes were placed into a temperature regulated gentleMACS Octo Dissociator™ (Miltenyi Biotech, Bergisch Gladbach, Germany) and underwent mechanical dissociation according to a custom mouse lung program at 37 ºC for 30 minutes. Dissociated tissue was passed through a pre-wetted 75 µm filter (Thermofischer Scientific, Burlington, ON, Canada), re-suspended with additional 5 mLs PBS and treated with 200 µLs 0.5M EDTA (Thermofischer Scientific, Burlington, ON, Canada). Pelleted cells were re-suspended in RBC-lysis buffer (Thermofischer Scientific, Burlington, ON, Canada) for 3 minutes at room temperature. Final cell pellet was re-suspended in 5 mLs PBS and cell counts and viability were performed using a Countess™ automated counter (Thermofischer Scientific, Burlington, ON, Canada).

**Flow cytometric analysis:** Single cell suspension was added to a v-bottom 96-well plate (Corning, NY, USA) 0.5 to $1x10^6$ cells/well. Live/Dead™ fixable staining assay was performed as instructed by product information (Thermofischer Scientific, Burlington, ON, Canada). Cells were blocked in FcR blocking reagent (Miltenyi Biotech, Bergisch Gladbach, Germany) for 15 min and incubated in 1:100 dilution PE-CD-144 (BD Biosciences, Mississauga, ON, Canada), PE-Dazzle™-CD-34 and APC-Fire™-CD-31 (BioLegend, San Diego, CA, USA) for 30 minutes. Cells were washed in FACS buffer (PBS, 1% BSA, 1mM EDTA) and fixed in 2% PFA for 10 minutes. 500 µLs of cells in FACS buffer were passed through a 40 µm mesh into flow tubes (BD Biosciences, Mississauga, ON, Canada). Samples were analyzed on a BD LSR Fortessa using BD FACSDIVA software for compensation and gating (Beckton Dickinson Biosciences, Franklin Lakes, NJ, USA). Further cytometric analysis were conducted on FlowJo v.10.6.2 (FlowJo LLC, Ashland, OR, USA).

**Immunohistochemistry**: Paraffin sections (3 µm thickness) were deparaffinized and rehydrated in a sequence of xylene, ethanol and $ddH_2O$ washes. Heat induced antigen retrieval with Citric Acid based antigen unmasking solution (Vector Labs, Burlingame, CA, USA) was performed. Sections were blocked in 10% FBS for 30 min and incubated with 1:400 goat anti-human HB-EGF (R&D Systems Inc., Oakville, ON, Canada) antibody over-night at 4 ºC. Following washes in PBS (0.25% Triton-X-100), sections were incubated with chicken anti-goat Alexa-594 secondary (Thermofischer Scientific, Burlington, ON, Canada) for 2 hours at room temperature. Images were taken on a Zeiss Axiophot epifluorescent microscope (Carl Zeiss Ltd., Toronto, ON, Canada) and processed with FIJI open source software (https://github.com/fiji).

**Evans blue assay**: A solution of 0.5% Evans blue (Eb) (Sigma-Aldrich, Oakville, ON, Canada) was prepared fresh in 0.9% sterile saline and passed through a 22 µm filter. 100 µL of Eb solution was injected via tail-vein and animals were allowed to rest for 2 hours before being sacrificed and bled. A catheter was inserted into the pulmonary artery and flushed with 10 mL of saline at a constant pressure (~25mmHg). Organs were collected and allowed to air dry for 2 hours. Dried tissue was weighted and place in centrifuge tubes with 500 µL of formamide (Sigma-Aldrich, Oakville, Ontario, Canada), incubated at 55 ºC for 48 hours and then centrifuged at 12,000 g for 20 minutes. Absorbance was measured in a plate reader at 610 nm wavelength. Concentration of Eb was cross referenced to standard curve and concentration per tissue weight was analyzed for each tissue and conditions collected.

**Micro-CT analysis:** Micro computed tomography (Micro-CT) was adapted to be used with mouse lungs from a previously published report (1).

**Hemodynamic measurements**: Animals were anaesthetized with ketamine (100 mg/kg) and xylazine (10 mg/kg) (i.p.) and placed into a 37 ºC heating pad. The right jugular vein was surgically exposed and a small incision was made with iris scissors. A mouse specific pressure sensor (Transsonic Systems Inc., London, ON, Canada) was inserted into vessel and placed into right ventricle (RV) for direct measurement of RVSP. Pressure loops were recorded and analyzed on LabScribe2 (Transsonic Systems Inc., London, ON, Canada). Right ventricular hypertrophy was determined by calculating the Fulton index, as defined by the weight ratio of the RV over left ventricle (LV) and septum (S), (RV/(LV+S)).

**MULTI-seq cell barcoding**: Barcoding of individual biological samples was performed as described by McGinnis *et.al.*(2019) (2). In summary, dissociated lung cells ($0.5x10^6$ cells per sample) were suspended in 150 µLs solution containing a 1:1 molar ratio (200nM) of anchor and barcode oligonucleotide containing a unique sequence for each of the 12 samples to be processed. Samples were incubated for 13 minutes at room temperature with gentle mixing every 3-5 minutes. Next, a co-anchor (200nM) was added to stabilize barcodes within membrane and incubated for additional 5 minutes. Cells were washed twice in PBS and cell counts were measured using a Countess$^{TM}$ automated counter (Thermofischer Scientific, Burlington, ON, Canada) and viability measured based on the ratio of cells staining with trypan blue (Thermofischer Scientific, Burlington, ON, Canada). Equal ratio of cells from each 12 barcodes were pooled and 1000 cells/µL were further processed through the 10x-Genomics pipeline. Only samples with viability > 85 % were used.

**Processing single-cell RNA sequencing libraries**: RNA library construction with 10x Genomics Single Cell 3’ RNA sequencing kit v3 was processed as previously described (3). Gene expression libraries were prepared as per manufacturer’s recommendations. Libraries for 48,347 cells were sequenced using a NextSeq500 (Illumina) with a mean of 8,550 reads per cell, and a median of 1,560 UMIs and 845 genes per cell. CellRanger v4.0 software (10x Genomics) was used to process raw sequencing reads with the mm10 reference transcriptome and with additional manual annotation of the diphtheria toxin receptor transgene. MULTI-seq barcode libraries were further trimmed to 28bp using Trimmomatic v.0.39 (https://github.com/timflutre/trimmomatic).

**Demultiplexing, doublet removal and quality control**: Barcodes were demultiplexed using the R package deMULTIplex (2) (https://github.com/chris-mcginnis-ucsf/MULTI-seq). Cells lacking barcodes or with multiple barcodes (doublets) were excluded from further analysis. Cells underwent an additional doublet removal step using the R package DoubletFinder (4) (https://github.com/chris-mcginnis-ucsf/DoubletFinder) and scDblFinder (https://github.com/plger/scDblFinder). Quality control was performed using R package Seurat (5) v.3.1.5 (https://github.com/satijalab/seurat). Cells with a high proportion (>30%) of mitochondrial transcripts and those with low complexity (<200 detected genes) were excluded from final matrix (Figure S4B and C). A total of 21,665 cells were used for downstream analyses. Data was log normalized and variable genes detected using “vst” method. Integrated analysis on top 3000 genes was conducted under “SCT” method where cell cycle and mitochondrial content and treatment condition were regressed out prior to calculation of PCA and UMAP on the first 40 principle components. Cell clusters were characterized using an automated annotation tool (6) and by cross referencing differential gene expression of individual clusters to previously characterized lung cells of the *Tabula Muris* cell atlas (7). The identity of the various lung cell clusters was further confirmed by the assessment of expression of cell-specific genes in the 21 clusters (Figure S5 and S6).

**Differential expression analysis:** Differential gene expression analysis was conducted using R package Muscat (8) multi-sample multi-group scRNA-seq analysis tools (https://github.com/HelenaLC/muscat). Standard workflow was performed by generating pseudobulk expression profiles for each cluster and testing for differential expression between experimental groups/conditions using default parameters unless otherwise specified.

**Automated cell classification:**
The different cell populations identified in our study were cross referenced with pre-annotated lung cells from the *Tabula Muris* (7) using the R package singleCellNet (https://github.com/pcahan1/singleCellNet).

**Cell type prioritization:**
To determine cells most affected during our different conditions in relation to control samples, we have employed a machine learning model to predict cells that become more separable during treatment based on their molecular measurements. For this we used the R package Augur (9) (https://github.com/neurorestore/Augur).

**Code availability:**
Code used to generate scRNAseq analysis will be provided at https://github.com/rsgodoy .

**Data availability:**
Data will be made available upon reasonable request made to the correspondence author.

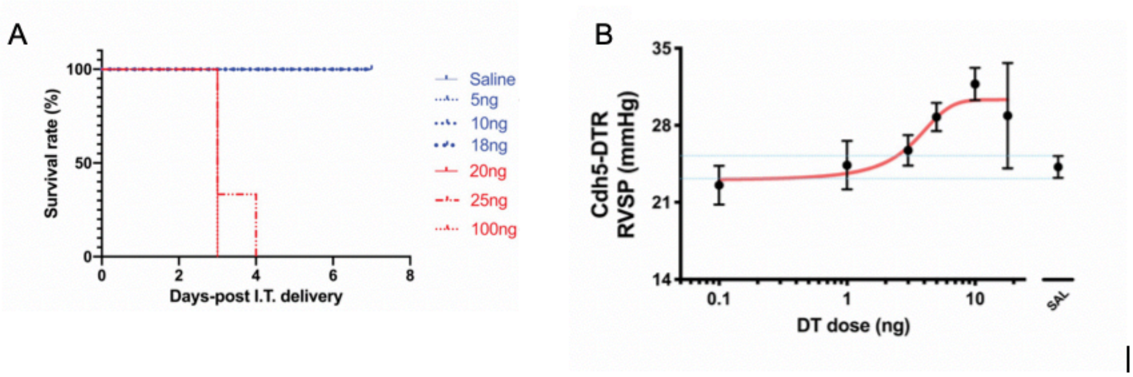


**Figure S1. Dose ranging studies for DT (0.1ng-18ng)**: A. Single intratracheal (IT) administration of DT at doses below 20ng were compatible with survival to 7 days post treatment. B. Doses above 1ng resulted in a dose-dependent increase in right ventricular systolic pressure (RVSP).

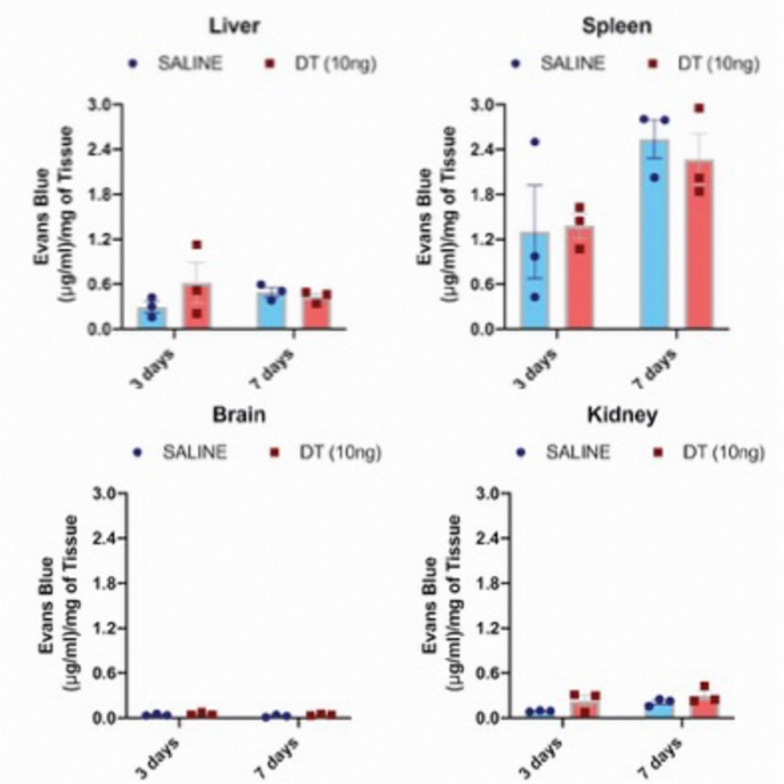


**Figure S2. Permeability is not increased in systemic vascular beds**: Vascular permeability as measured by Evans blue uptake remained unaltered following DT administration at 3 and 7 days in liver, spleen, kidney and brain.

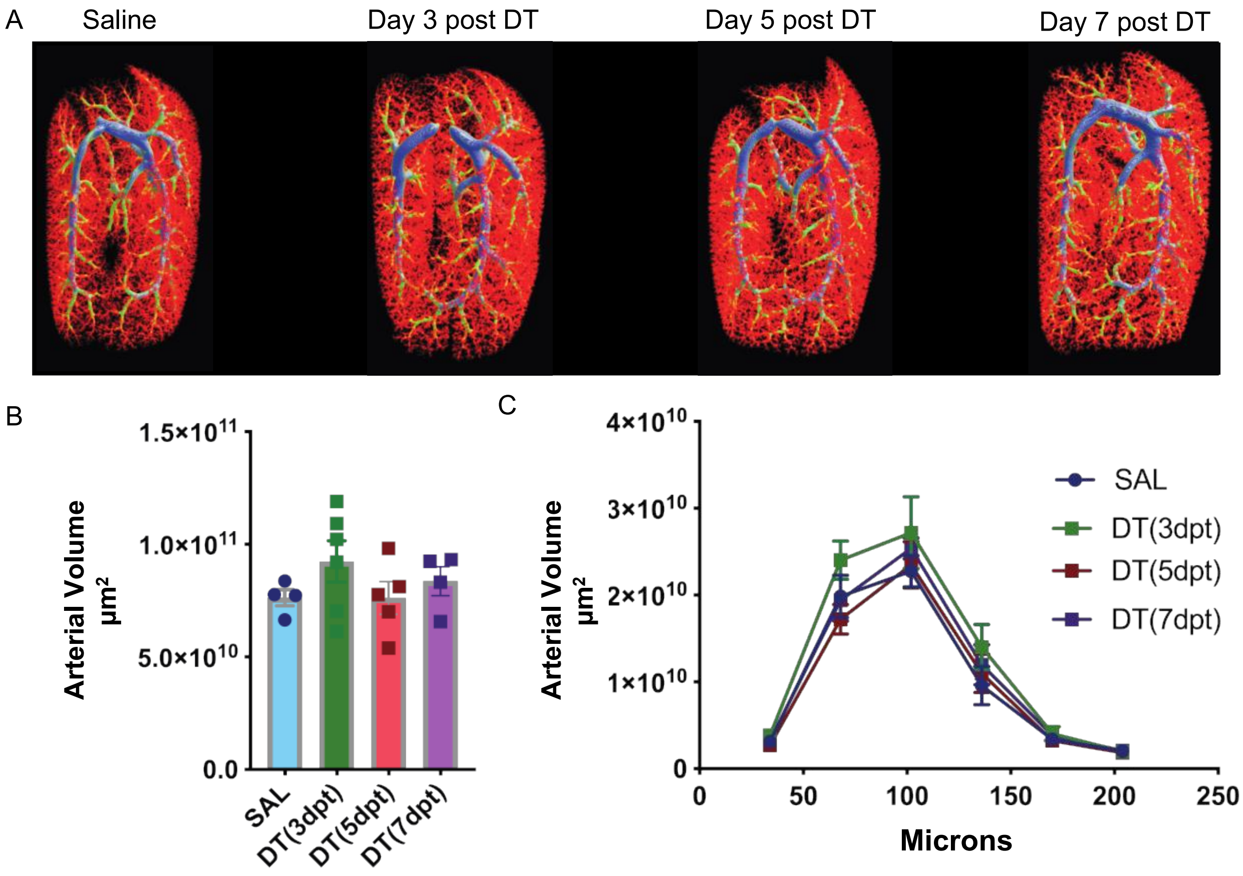


**Figure S3. Lack of distal arteriolar pruning in the DT-induced model of ALI**: A) Representative examples of high-resolution micro-CT images in saline control mice and 3, 5 and 7 days post DT-induced EC ablation. B) Summary data showing no change in arterial volumes at the different timepoints post DT compared with saline controls. C) Distribution of arterial volumes based on arterial size. There was no change in the size distribution of arterial volume between groups with the greatest volumes in arterioles < 200 microns. dpt = days post DT treatment.

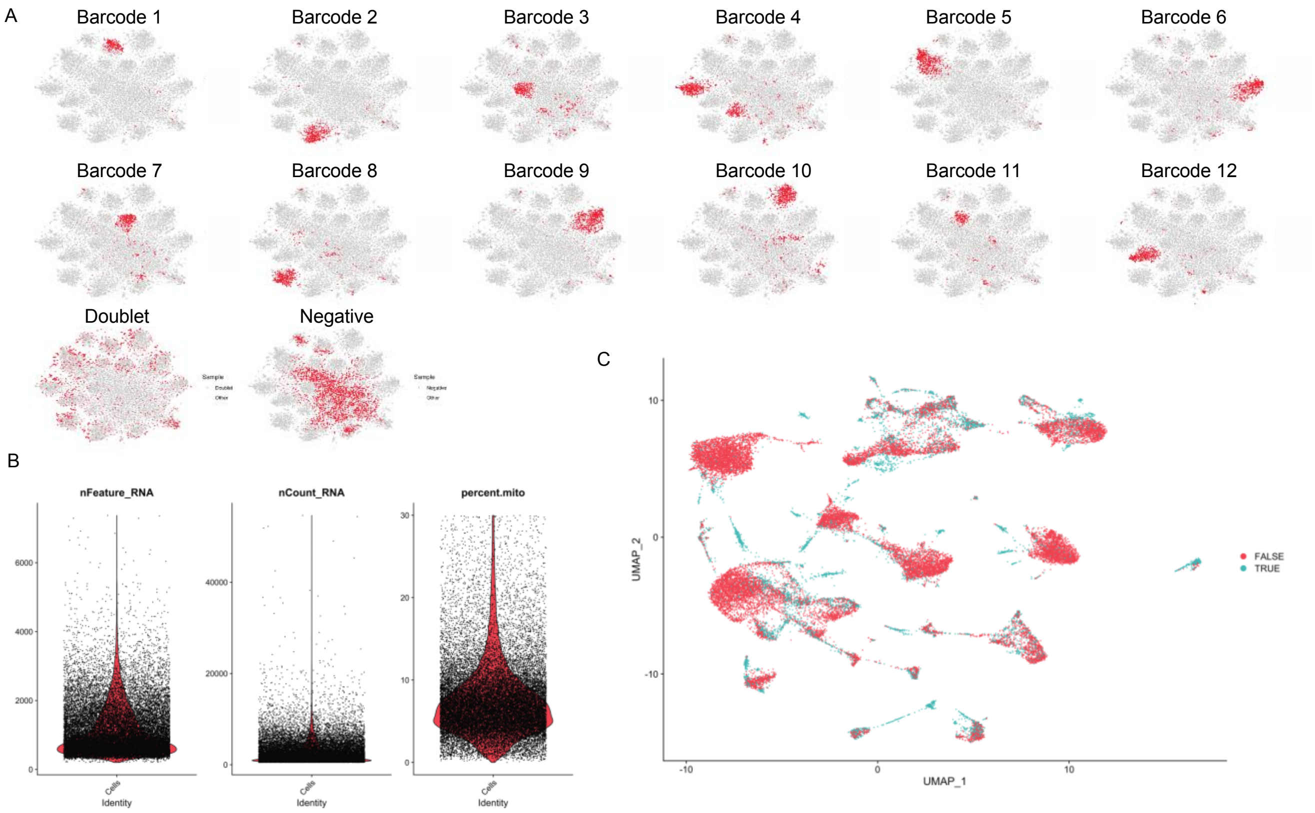


**Figure S4. Quality control for scRNAseq**: A) All 12 barcodes were demultiplexed with a small number of cells with multiple barcodes (doublets) and a population lacking any barcode (negative). B) Quality control was performed using the number of features, counts and percentage of mitochondrial content as cutoff. C) A further computational method for doublet removal was conducted in addition to excluding doublets based on multiple barcode uptake.

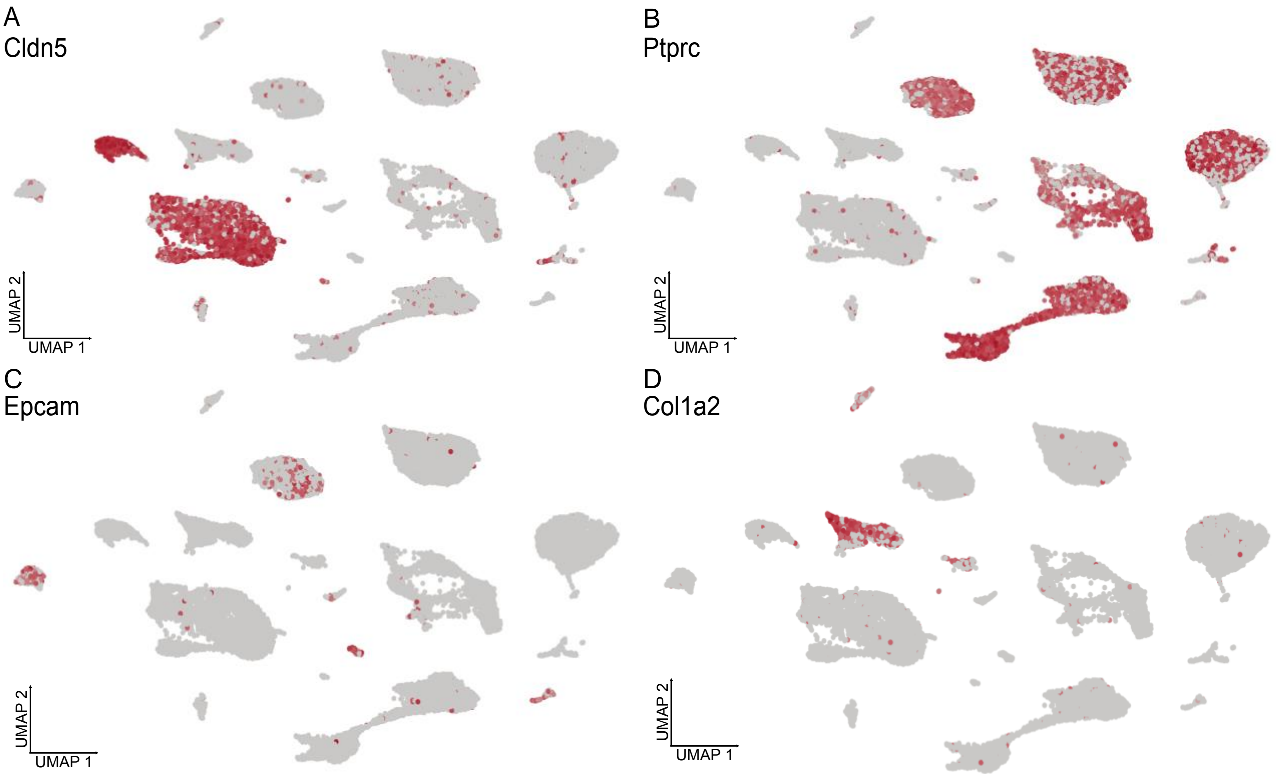


**Figure S5. Major cell type classification:** Expression of selected genes characterizing the identity of the major lung cell clusters based on the *Tabula Muris* biological atlas, including *Cldn5* (endothelial) (A), *Ptprc* (immune) (B), *Epcam* (epithelial) (C), and *Col1a* (mesenchymal) (D).

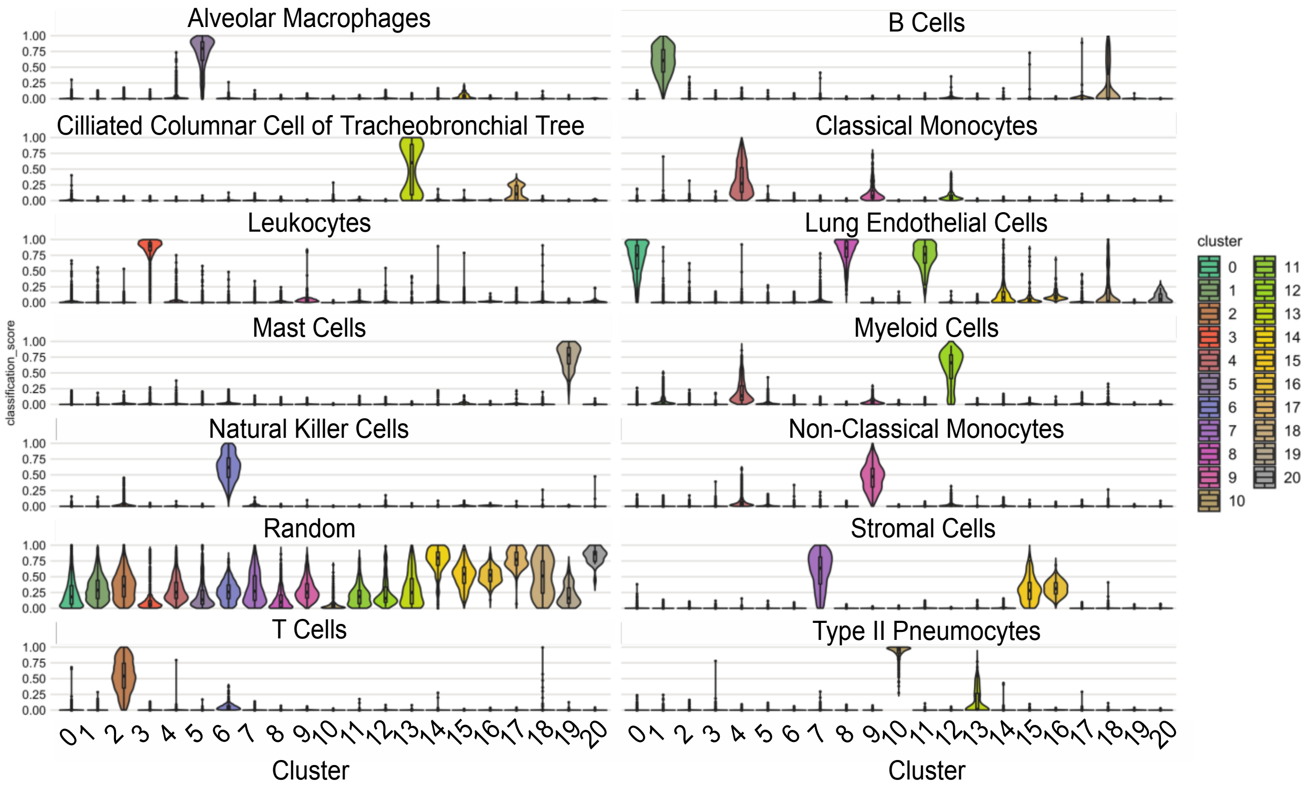


**Figure S6. Automated cell classification:** Violin plots of cell-specific gene expression in the major lung cell clusters based on the *Tabula Muris* biological atlas.

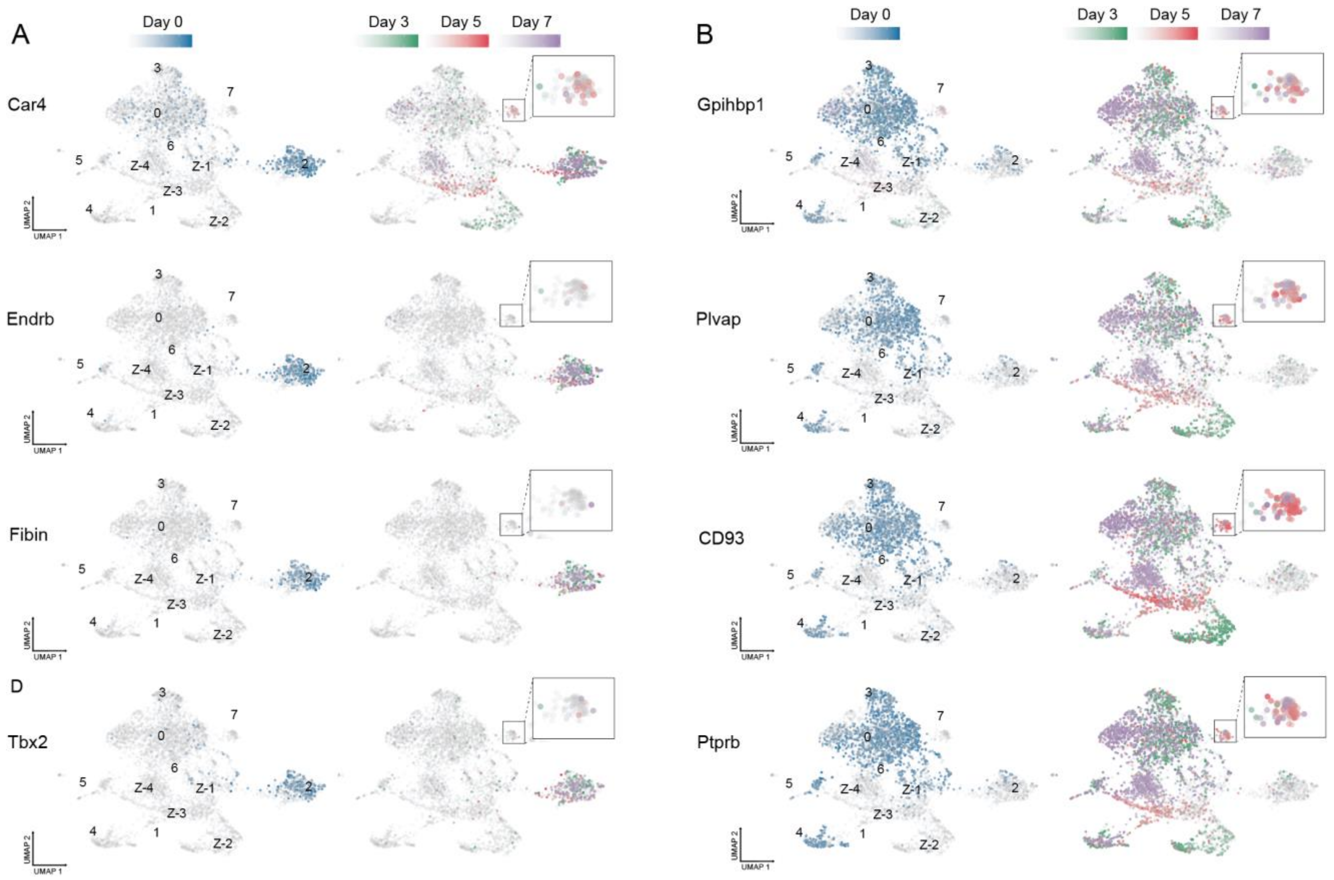


**Figure S7. aCap and gCap genes in EC populations pre and post injury**: A) Typical aCap ECs genes include: *Car4* (Carbonic Anhydrase 4); *Endrb* (endothelin receptor type-B); *Fibin* (Fin Bud Initiation Factor Homolog); and *Tbx2* (T-Box Transcription Factor 2). B) In contrast, gCap genes include: *Gpihbp1* (Glycosylphosphatidylinositol Anchored High Density Lipoprotein Binding Protein 1); *Plvap* (Plasmalemma Vesicle Associated Protein); *Cd93*; and *Ptprb* (Protein Tyrosine Phosphatase Receptor Type B). Day 3 (green); Day 5 (red) and Day 7 (purple) with cluster 7 enlarged in the insets.

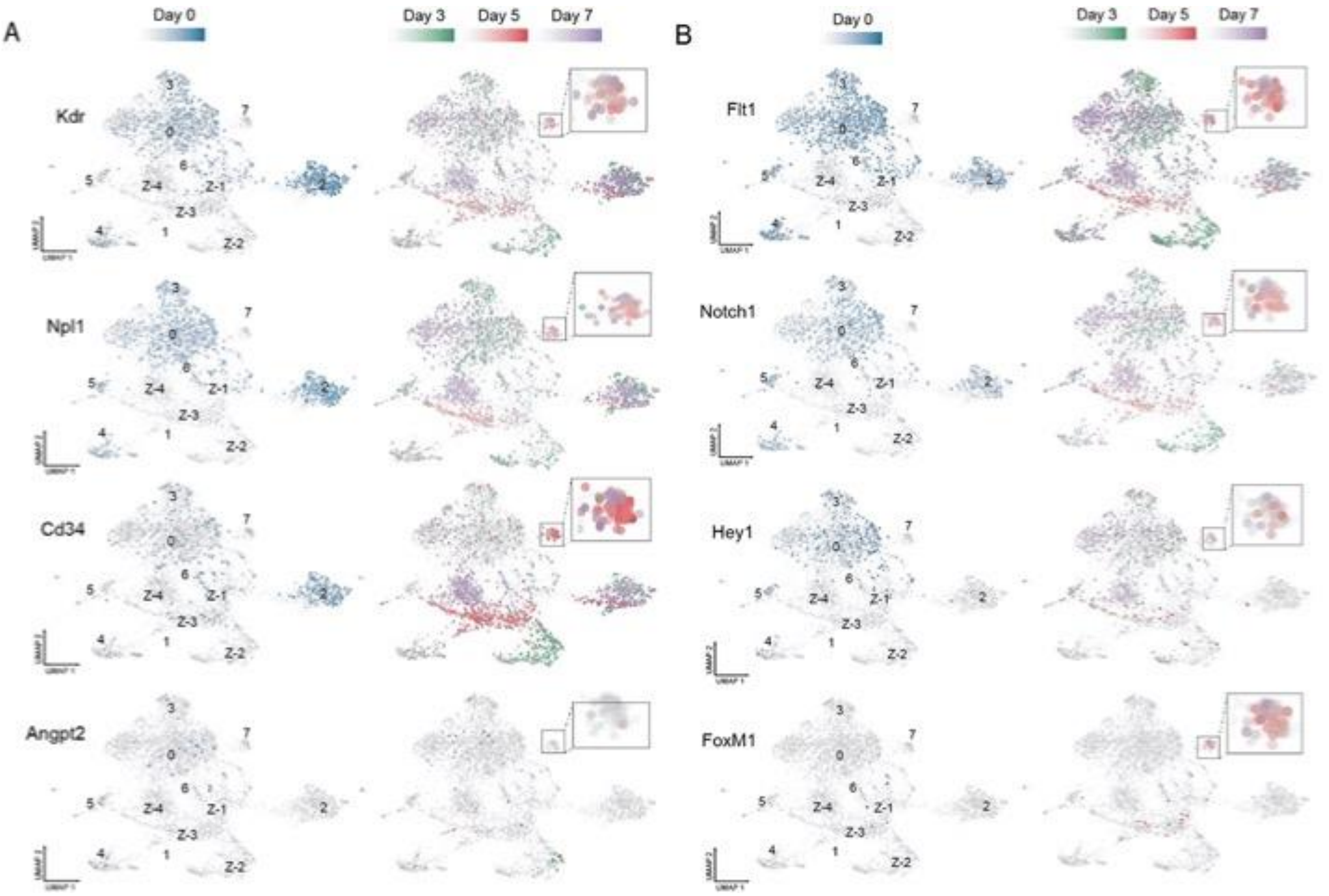


**Figure S8. Temporal evolution of expression of angiogenic genes pre and post EC ablation**: A) *Kdr* (VEGFR2), *Npl1* (Neuropilin-q), *Cd34* and *Angpt2* (Angipoopietin-2). B) *Flt1* (VEGFR1), *Notch1*, *Hey1* and *FoxM1*. Lung ECs from control mice (blue) are shown on the left and cells from the DT-treated cohorts are presented on the right: Day 3 (green); Day 5 (red) and Day 7 (purple) with cluster 7 enlarged in the insets.

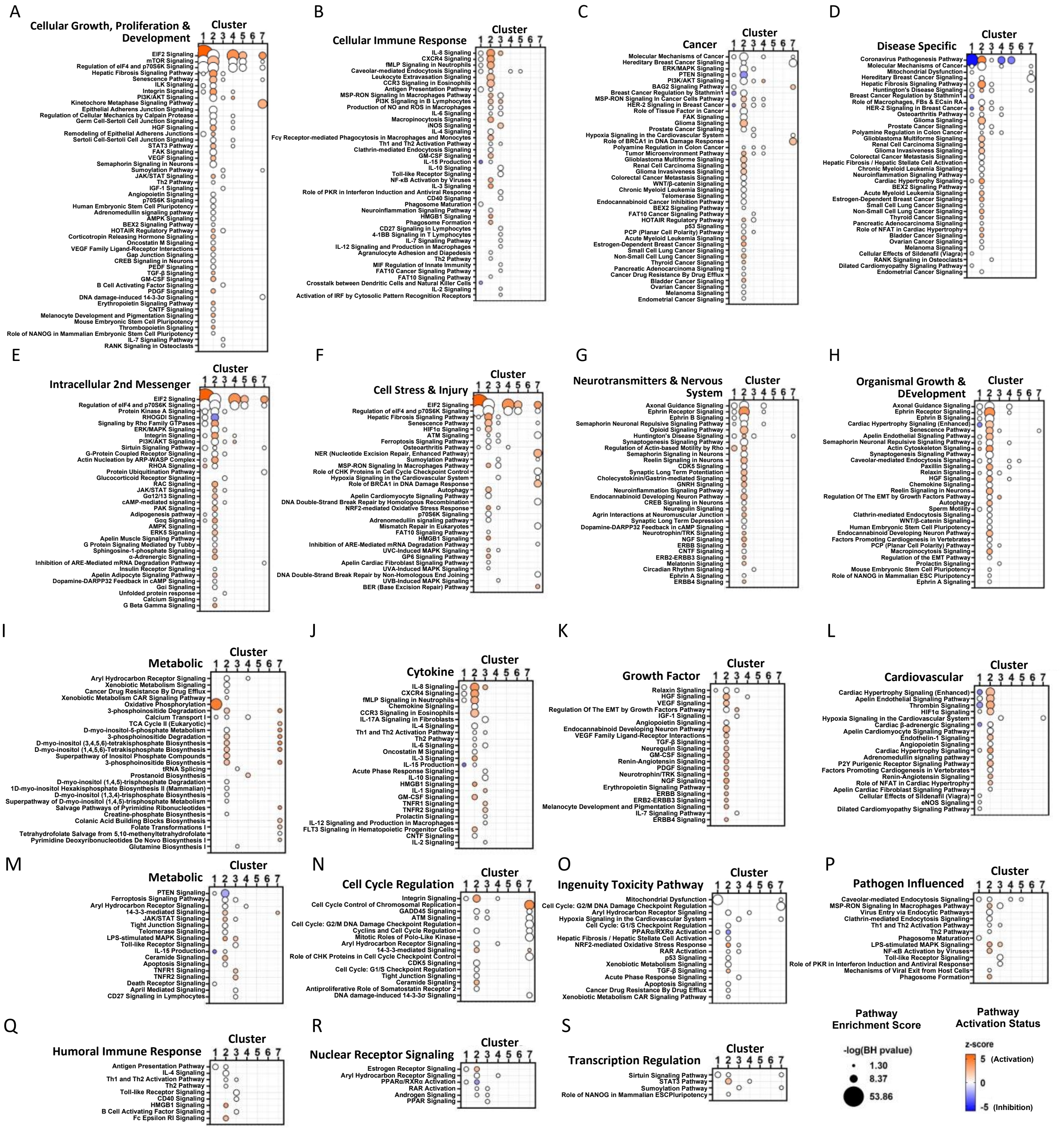


**Figure S9**. **Pathway enrichment analysis of endothelial clusters 1 to 7 via Ingenuity Pathway Analysis**. Each panel shows a predefined functional category and specific canonical signaling pathways that were significantly enriched in genes altered in each endothelial cluster 1 to 7 (x-axis). Circle size denotes pathway enrichment score based on the Benjamini-Hochberg (BH) corrected p-value. A minimum score of 1.3 (i.e., FDR<0.05) was used as an inclusion threshold. Z-score color denotes the predicted activation state of the pathway based on the degree of matching between the expected and observed pattern of gene expression changes. Only pathways with z-score >+2 (activated) or <-2 (inhibited) are shown in color to highlight confident predictions. White circles denote pathways in which the activation state cannot be confidently predicted. No pathways were significantly enriched in DE genes from Cluster 6 at the FDR<0.05 threshold.

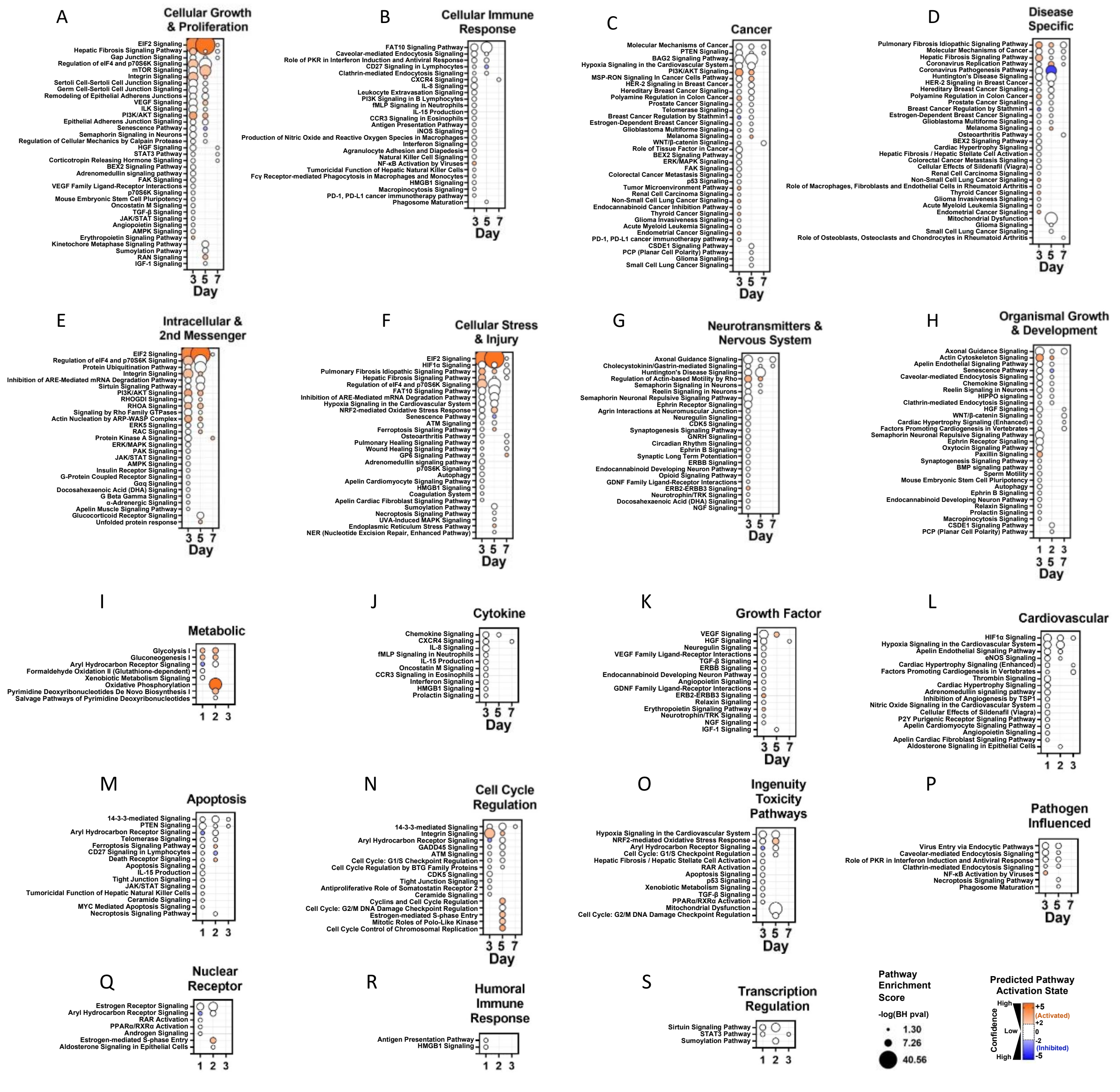

**Figure S10. Pathway enrichment analysis of time-dependent changes in endothelial cluster 1 at days 3, 5 and 7 versus day 0**. Each panel shows a predefined functional category and specific canonical signaling pathways (y-axis) that were significantly enriched in genes altered at each time point (days 3, 5 and 7 versus day 0) in endothelial cluster 1 (x-axis). Circle size denotes pathway enrichment score based on the Benjamini-Hochberg (BH) corrected p-value. A minimum score of 1.3 (i.e., FDR<0.05) was used as an inclusion threshold. Z-score color denotes the predicted activation state of the pathway based on the degree of matching between the expected and observed pattern of gene expression changes. Only pathways with z-score >+2 (activated) or <-2 (inhibited) are shown in color to highlight confident predictions. White circles denote pathways in which the activation state cannot be confidently predicted. Of note, some canonical pathways may be classified in more than one category.
